# Cell-type-specific plasticity in synaptic, intrinsic, and sound response properties of deep-layer auditory cortical neurons after noise trauma

**DOI:** 10.1101/2025.07.09.663954

**Authors:** Yanjun Zhao, Stylianos Kouvaros, Nathan A. Schneider, Rebecca F. Krall, Stephanie Lam, Megan P. Arnold, Ross S. Williamson, Thanos Tzounopoulos

**Affiliations:** Department of Otolaryngology, University of Pittsburgh, Pittsburgh, PA; Department of Neurobiology, University of Pittsburgh, Pittsburgh, PA; Department of Bioengineering, University of Pittsburgh, Pittsburgh PA; Pittsburgh Hearing Research Center, University of Pittsburgh, Pittsburgh, PA; Center for the Neural Basis of Cognition, University of Pittsburgh, Pittsburgh, PA; Center for Neuroscience, University of Pittsburgh, Pittsburgh, PA; Department of Biomedicine, University of Basel, Basel, CH

**Keywords:** auditory cortex, noise-induced hearing loss, plasticity, thalamocortical circuits, excitatory neurons, in vivo imaging, in vitro electrophysiology

## Abstract

Peripheral trauma, such as noise-induced hearing loss (NIHL), triggers compensatory plasticity in the auditory cortex (ACtx) to maintain auditory function. While cortical plasticity in superficial cortical layers has been relatively well studied, the plasticity mechanisms governing deep-layer excitatory projection neurons remain less understood. Here, we investigated the plasticity of layer (L)5 extratelencephalic (ETs) and L6 corticothalamic neurons (CTs) following NIHL. Using a combination of *in vitro* slice electrophysiology, optogenetics, and *in vivo* two-photon imaging in a mouse model of NIHL, we characterized changes in evoked thalamocortical (TC) synaptic input strength, intrinsic excitability, and sound response properties. We found that TC input was initially equivalent between ETs and CTs, then shifted to CT-dominant one day after noise exposure. This shift renormalized to equivalent seven days after noise exposure and was associated with a transient increase in both the quantal size (*q*) in TC→CT synapses and intrinsic CT suprathreshold excitability. ETs maintained stable intrinsic properties and showed minor changes in their TC input. *In vivo* imaging revealed that CTs displayed a persistent elevation in sound intensity thresholds, whereas ETs transiently shifted their best frequency representation and reduced their responsiveness to high-frequency tones one day after NIHL, followed by recovery at seven days. Together, our findings highlight cell-type-specific plasticity mechanisms in deep-layer cortical neurons, enhance our understanding of cortical adaptation to peripheral damage, and highlight targets for developing therapeutic strategies to mitigate hearing loss and related disorders such as tinnitus and hyperacusis.

**Short Abstract:** Peripheral damage drives auditory cortex (ACtx) plasticity, but the underlying synaptic and cellular mechanisms remain poorly understood. We used a combination of *in vitro* slice electrophysiology, optogenetics, and *in vivo* two-photon imaging to investigate layer 5 extratelencephalic (ET) and layer 6 corticothalamic (CT) neuronal plasticity in mice, following noise-induced hearing loss (NIHL). Thalamocortical (TC) input was initially balanced between CTs and ETs but shifted to CT-dominant one day post-NIHL and then normalized by day seven. This transient shift was accompanied by increased quantal size and suprathreshold excitability in CTs, with minimal changes in ETs. *In vivo*, CTs exhibited persistent elevation in sound intensity thresholds, while ETs showed a transient shift in frequency tuning and reduced high-frequency responsiveness that recovered within a week. These findings reveal distinct, cell-type-specific plasticity mechanisms in deep-layer ACtx neurons following peripheral damage and highlight potential targets for treating hearing loss-related disorders such as tinnitus and hyperacusis.

## Introduction

In all sensory systems, damage to peripheral organs induces compensatory plasticity mechanisms in the brain, which contributes to function preservation by increasing neural responsiveness to surviving sensory inputs. In the cortex, this homeostatic plasticity is characterized by increased central gain and has been well documented across sensory modalities (Rasmusson, 1982; Merzenich et al., 1983; Robertson and Irvine, 1989; Kaas et al., 1990; Gilbert and Wiesel, 1992). In the auditory system, exposure to loud noise induces cochlear damage that often leads to a reduction in auditory nerve input due to cochlear synaptopathy and the degeneration or death of both hair and spiral ganglion cells (Ryan and Bone, 1978; Liberman and Dodds, 1984; Kujawa and Liberman, 2009). Despite this peripheral damage, the sound-evoked activity of excitatory cortical principal neurons (PNs) is frequently restored, or even enhanced, within days of NIHL (Qiu et al., 2000; Auerbach et al., 2014; Chambers et al., 2016; Resnik and Polley, 2017; Kumar et al., 2023), reflecting robust cortical and subcortical homeostatic plasticity mechanisms. These mechanisms support perceptual hearing threshold recovery, especially in quiet backgrounds, even after a dramatic loss of peripheral input (Chambers et al., 2016; Resnik and Polley, 2021)

In the ACtx, noise-induced cochlear damage is associated with enhanced central gain, reduced cortical inhibitory (GABAergic) activity, increased spontaneous firing, and reorganization of frequency tuning towards less damaged regions of the cochlea (Seki and Eggermont, 2003; Kotak et al., 2005; Sarro et al., 2008; Scholl and Wehr, 2008; Yang et al., 2011; Takesian et al., 2013; Chambers et al., 2016; Resnik and Polley, 2017). Recent studies in ACtx layer (L) 2/3 neurons revealed that this plasticity is mediated by cell-type-specific mechanisms enabling a precise division of labor and cooperativity amongst cortical interneurons (Kumar et al., 2023). However, the plasticity mechanisms governing excitatory projection neurons remain less understood. Addressing this knowledge gap requires examining the distinct contributions of deep-layer neurons, specifically L5 extratelencephalic neurons (ETs, also known as pyramidal tract or PT neurons) and L6 corticothalamic neurons (CTs), as L5 and 6 represent the primary output pathways from the ACtx. CTs exclusively target the thalamus and readily modulate local cortical gain, while also modulating ongoing thalamocortical oscillatory activity, forming key nodes in cortico-thalamo-cortical loops (Olsen et al., 2012; Guo et al., 2017; Ibrahim et al., 2021; Shepherd and Yamawaki, 2021). ETs, in addition to the thalamus, also project to the midbrain, striatum, and amygdala (Harris and Shepherd, 2015; Chen et al., 2019; Williamson and Polley, 2019; Shepherd and Yamawaki, 2021; Issa et al., 2023). These projections are crucial for connecting the auditory system to downstream targets involved in motor control, sensorimotor integration, learning, and higher-order cognitive processes (Winer, 2005; Bajo et al., 2010; Stebbings et al., 2014; Xiong et al., 2015; Ford et al., 2024; Quass et al., 2024). It is therefore essential to determine how ETs, CTs, as well as their thalamocortical inputs change after NIHL. Given the importance of cortico-thalamo-cortical loops in perception and cognition in normal and pathological states (Tononi and Edelman, 1998; Tononi et al., 1998; Jones, 2001; Llinas et al., 2005; Sherman, 2007; Shepherd and Yamawaki, 2021), understanding these mechanisms may highlight novel approaches to enhance perceptual recovery after hearing loss and/or mitigate plasticity-induced disorders such as tinnitus and hyperacusis (Henton and Tzounopoulos, 2021).

To address these gaps in knowledge, we employed a mouse model of NIHL to investigate plasticity mechanisms after noise trauma. We used a combination of *in vitro* slice electrophysiology and *in vivo* two-photon imaging to characterize changes in thalamocortical synaptic strength to ETs and CTs, and intrinsic and sound response properties of ETs and CTs after NIHL. Our findings highlight cell-type-specific plasticity mechanisms, offering insights into cortical plasticity in response to peripheral damage.

## Materials and Methods

### Animals

All mice handling was approved by the Institutional Animal Care and Use Committee at the University of Pittsburgh. 66 male and 38 female Ntsr1-Cre mice (MMRRC, B6.FVB(Cg)-Tg(Ntsr1-Cre)GN220Gsat/Mmcd), were used for experiments shown in Figures 1-5, S1-2; 7 male and 2 female C57Bl/6J mice were used for Figure 6; 3 male and 1 female Ntsr1-Cre mice as well as 4 male and 5 female Ntsr1-Cre x Ai148 mice (JAX, Ai148(TIT2L-GC6f-ICL-tTA2)-D) were used for Figure 7.

**Figure 1.**
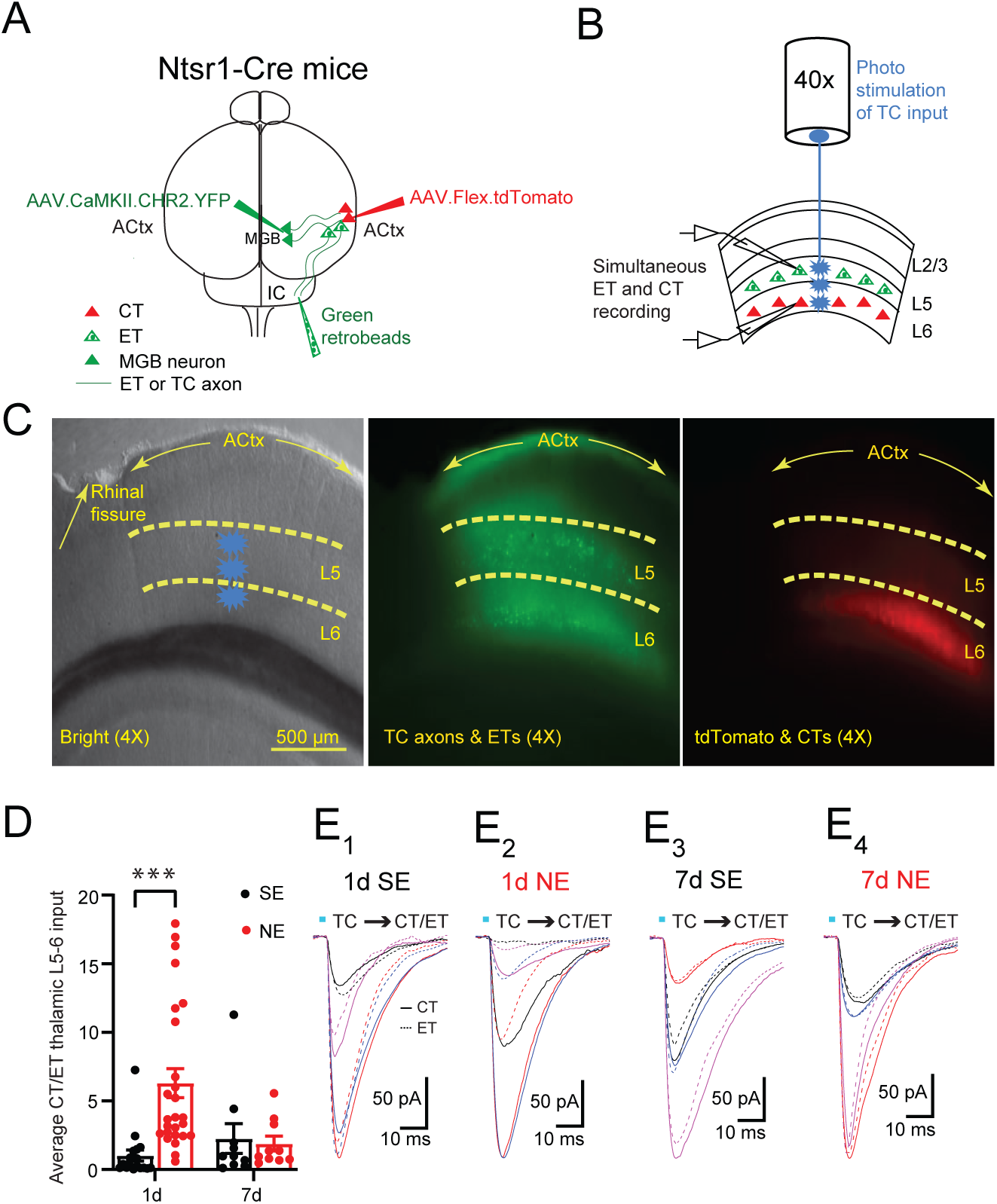
Transient shift in TC synaptic strength from CT and ET equivalent to CT dominant one day after NE. **(A)** Schematic illustration of stereotaxic injections of retrograde microspheres to label L5 extratelencephalic neurons (ETs, green), and viral vectors (AAVs) for expression of ChR2 in thalamocortical (TC) inputs and tdTomato in L6 corticothalamic neurons (CTs, red) in Ntsr1-Cre mice. **(B)** Schematic illustration of slice electrophysiology experiment involving photostimulation of ChR2 expressing TC afferents and simultaneous (dual) recording from an ET (green) and a CT (red). **(C)** Images in 4X magnification showing the extent of ACtx area in bright-field (left), green-labeled ETs and TC axons (middle) and red-labeled CTs **(D)** Average CT/ET EPSC ratio after optogenetically stimulating thalamic L5-6 input 1d and 7d after SE and NE. (1d SE: 18 cells/6 mice; 1d NE: 26 cells/9 mice; 7d SE: 10 cells/6 mice; 7d NE: 10 cells/6 mice). Asterisks indicate significant differences (***p<0.001, two-way ANOVA and Bonferroni correction for multiple comparisons). **(E)** Representative traces of EPSCs in dual recordings from both CT (solid line) and ET (dotted line) neurons evoked by maximal photostimulation of L5-6 thalamocortical inputs in 1d SE (E_1_) and 1d NE (E_2_); 7d SE (E_3_) and 7d NE (E_4_). Different colored traces represent different pairs of simultaneously recorded CTs and ETs. Detailed statistical values are listed in Table 1.

**Table 1.**
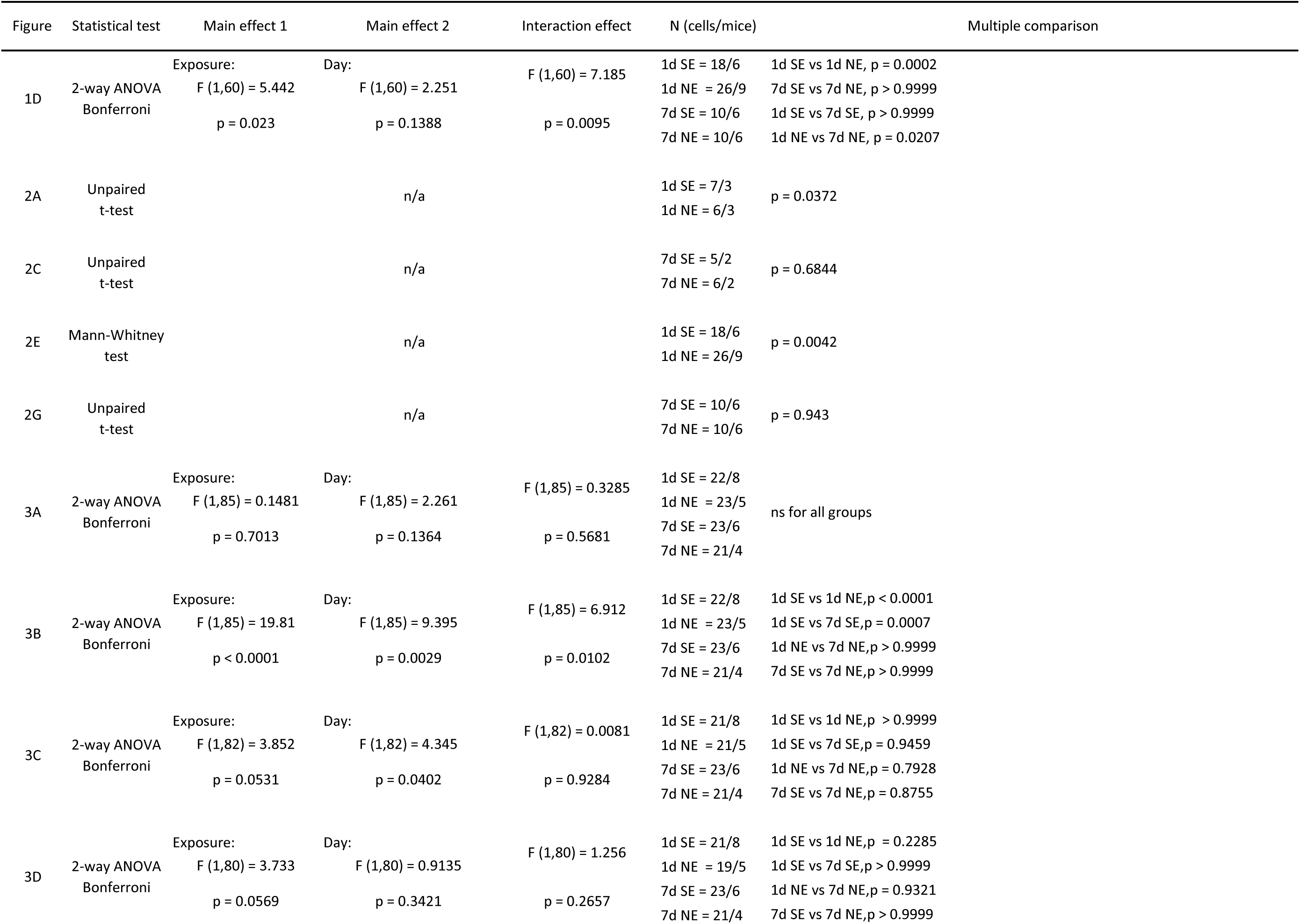
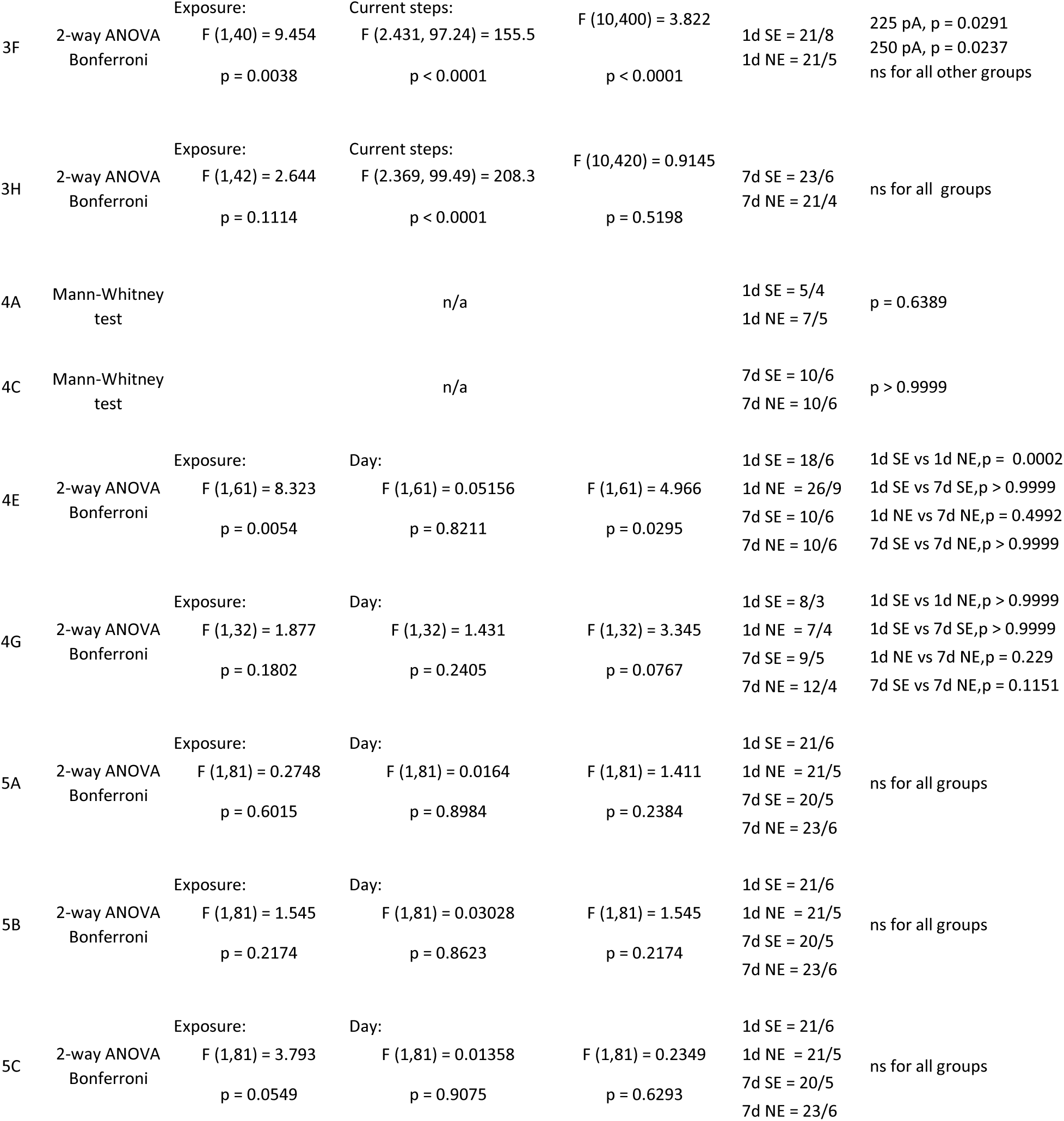
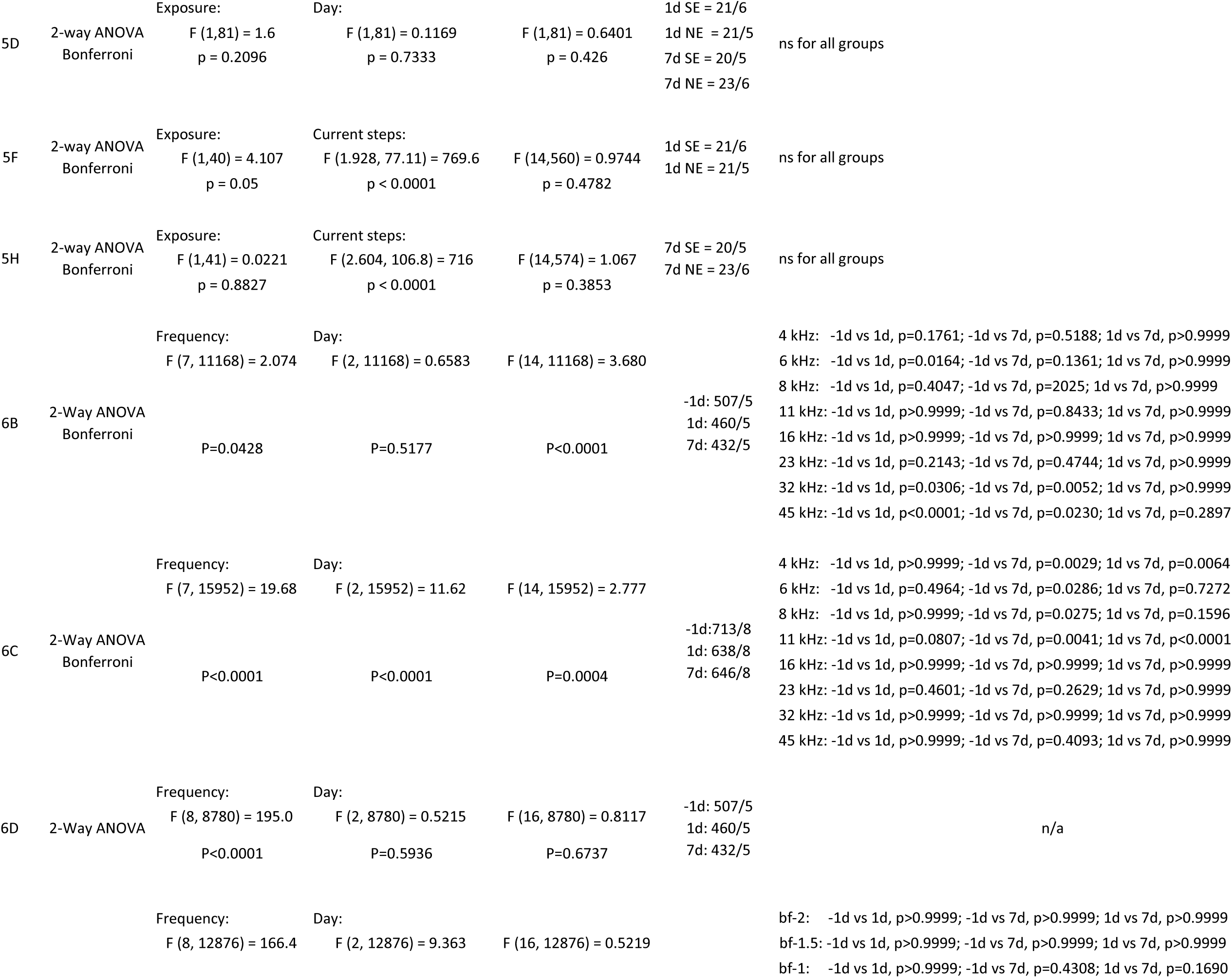
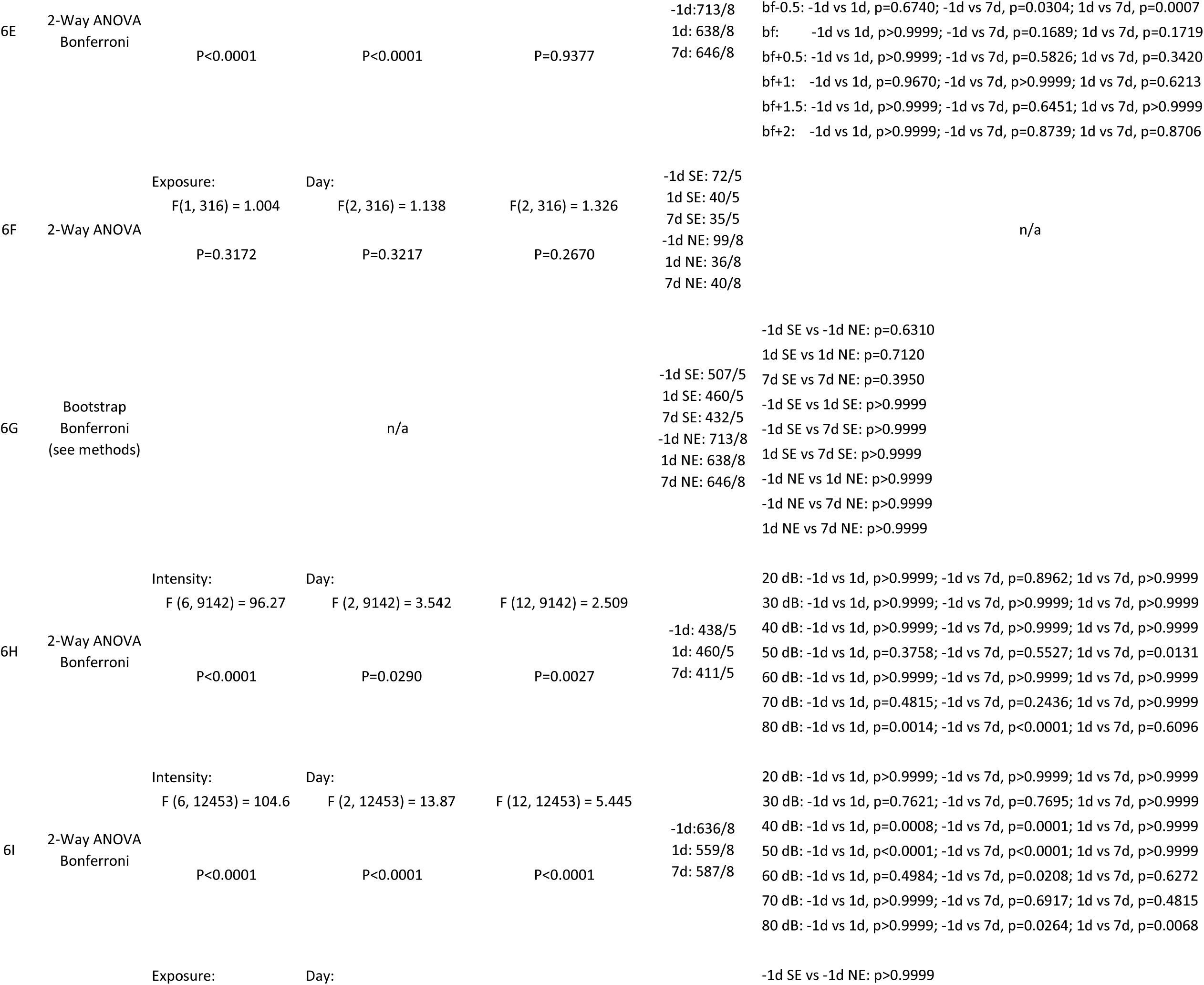
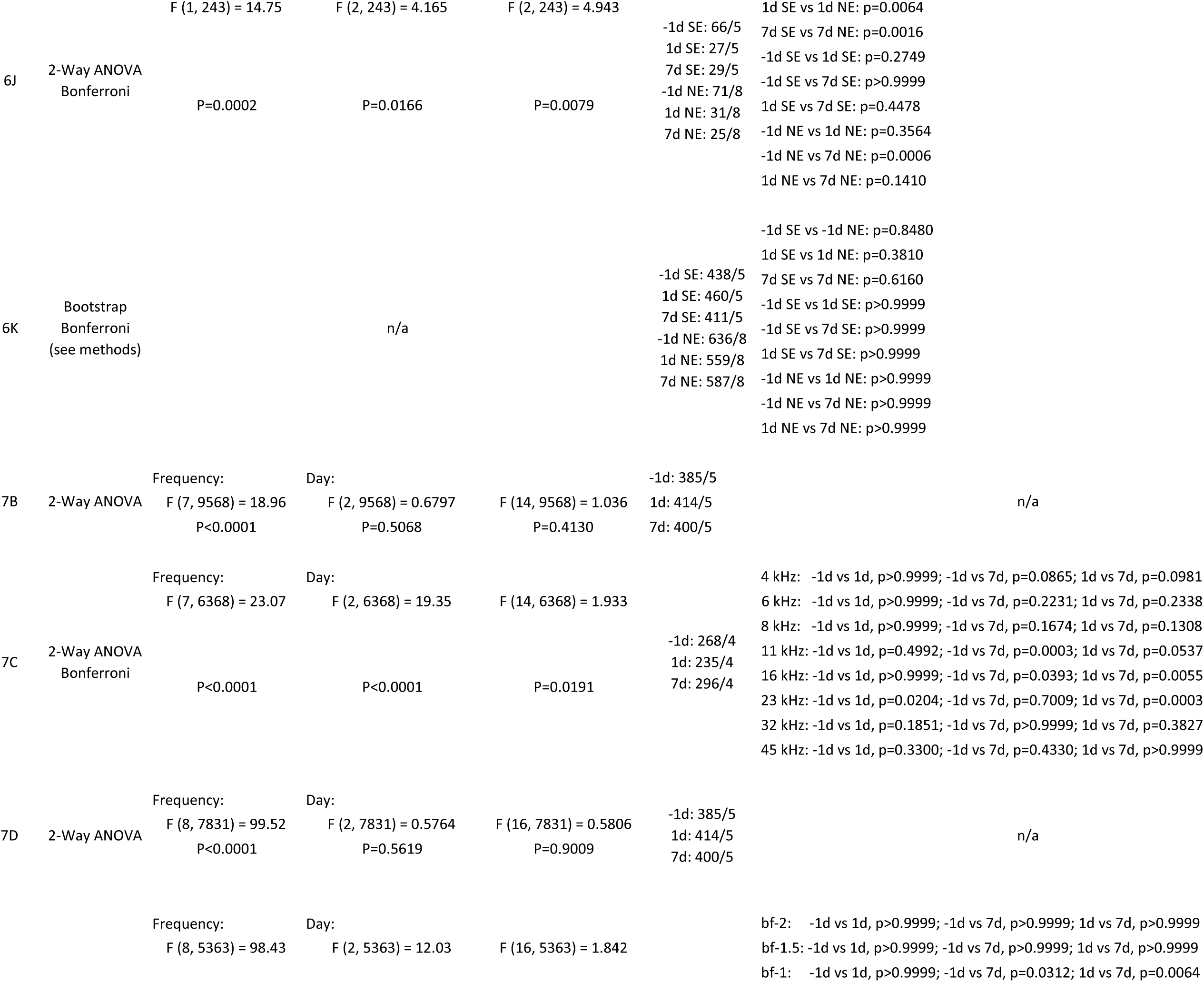
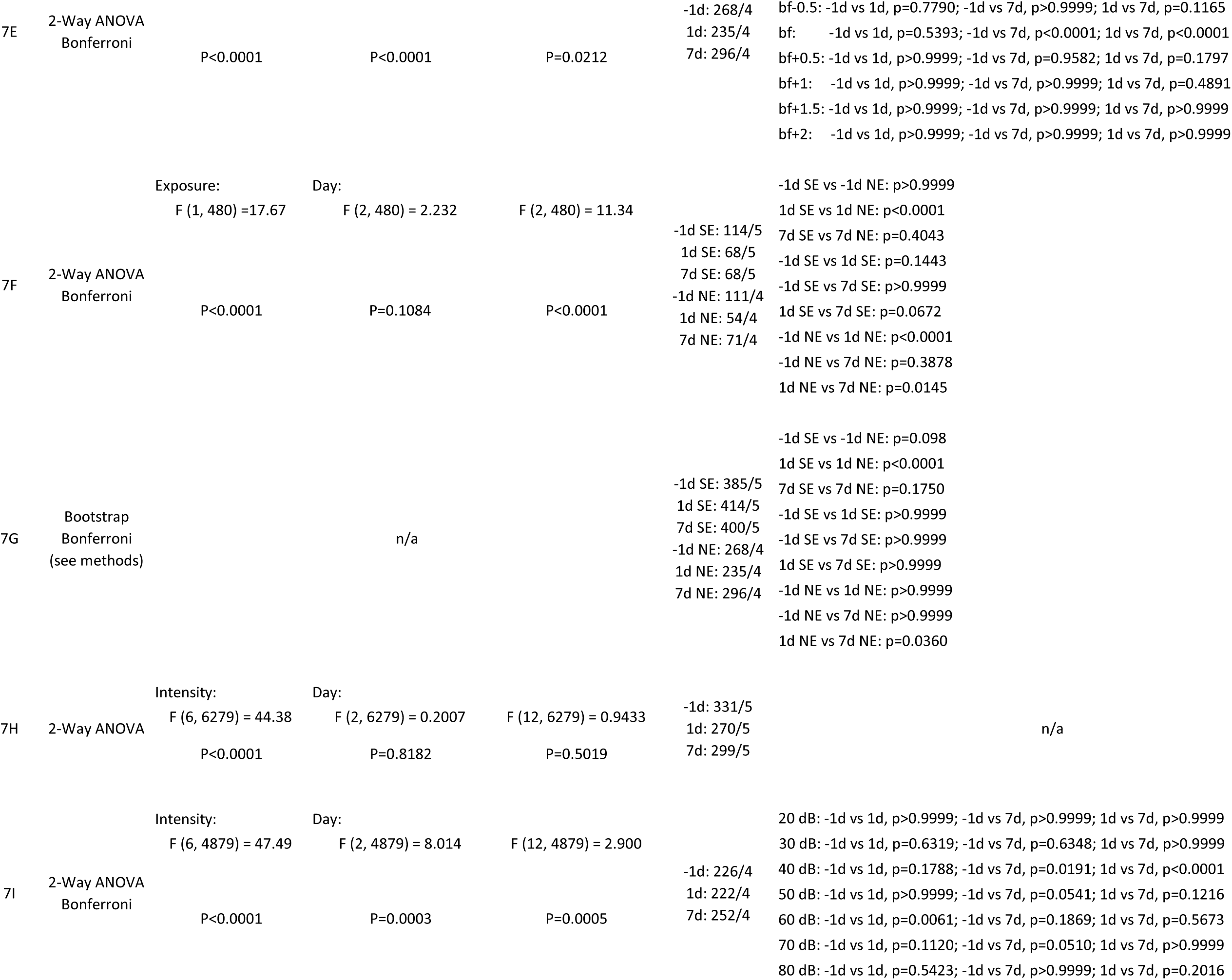
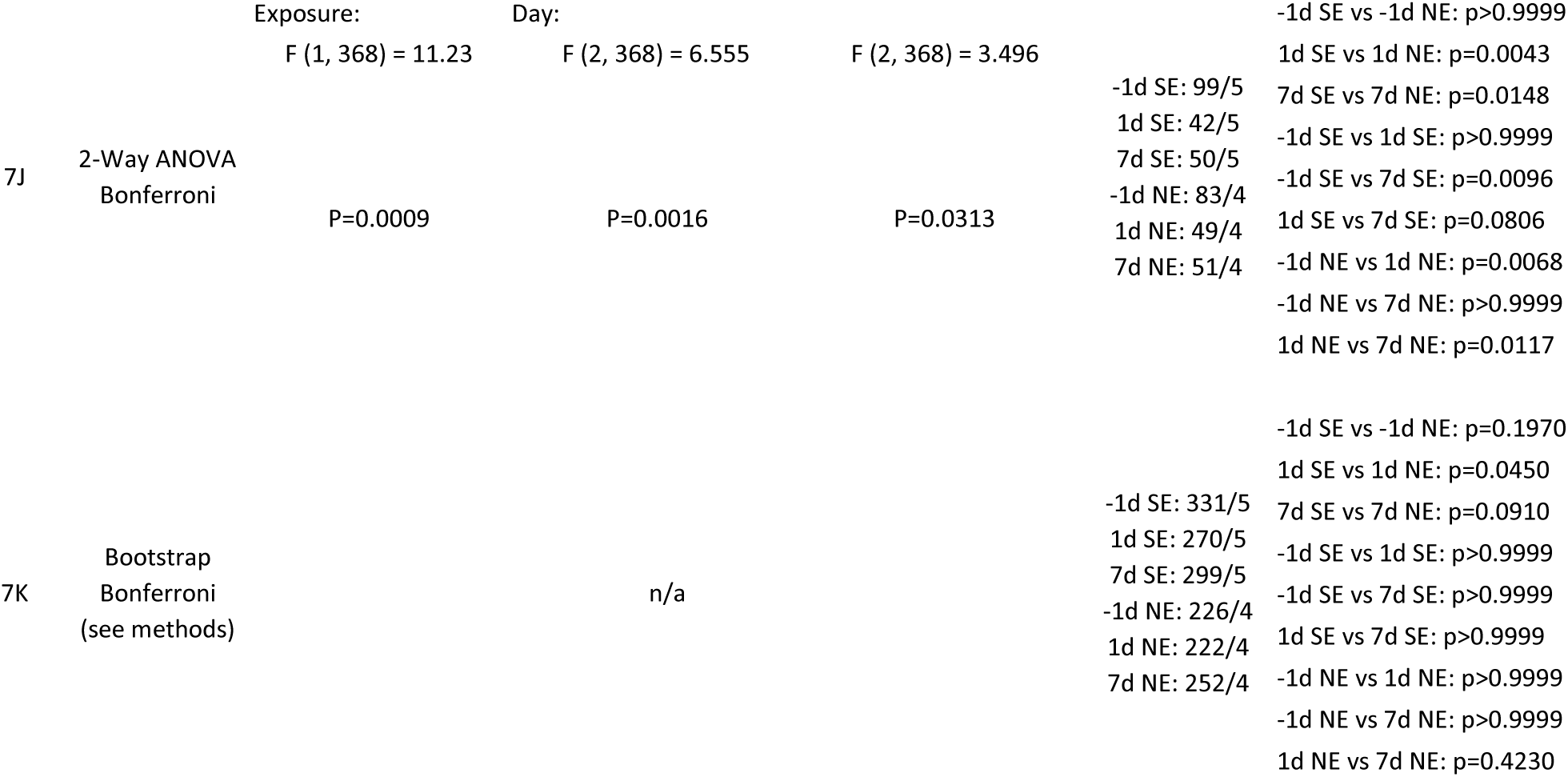
Statistical values for Figures 1-7.

### Surgical methods for *in vitro* electrophysiology

Mice (P28-35) were anesthetized with isoflurane (induction: 3% in oxygen, maintenance: 1.5-2% in oxygen) and secured in a stereotaxic frame (Kopf). Core body temperature was maintained at ∼37°C with a heating pad and eyes were protected with ophthalmic ointment. Lidocaine (1%) was injected under the scalp and a midline incision was made to expose the skull. ETs were retrogradely labeled by injecting green colored fluorescent latex microspheres (Lumafluor) into the right inferior colliculus (1 mm posterior to lambda and 1 mm lateral, at an injection depth of 0.75 mm), as schematized in **Figure 1A**. To transduce CTs, an AAV (AAV9-FLEX-tdTomato, titer: 1e^13^ gc/mL; Addgene) was injected into the right ACtx of Ntsr1-Cre mice through a small burr hole (∼0.4 mm diameter, ∼4 mm lateral to lambda). A glass micropipette containing the viral vector was lowered into ACtx to a depth of ∼1 mm using a micromanipulator (Kopf). A syringe pump (World Precision Instruments) was then used to inject 200 nL (diluted 1:1 in PBS) over the course of 2 minutes, at a rate of 100 nL/min.

To facilitate stimulation of thalamic afferents, 500 nL (undiluted) of AAV9-CaMKIIa-hChR2(H134R)-EYFP (titer: 8.96e^13^ gc/ml, Addgene) was injected into the right medial geniculate body of Ntsr1-Cre mice (3 mm posterior/2 mm lateral/3mm depth) at a rate of 100 nL/min, alongside injections of microspheres and AAV-FLEX-tdTomato, as described earlier. Post injection, the scalp was closed with cyanoacrylate adhesive and mice were administered with a subcutaneous injection of carprofen (5 mg/kg).

### Surgical methods for *in vivo* two-photon imaging

Mice (P56-72) were anesthetized with isoflurane (induction: 5% in oxygen, maintenance: 1.5-2% in oxygen) and secured in a stereotaxic frame (Kopf). Dexamethasone (2 mg/kg) was administered via an intraperitoneal injection to mitigate brain edema. Core body temperature was maintained at ∼37°C with a heating pad and eyes were protected with ophthalmic ointment. Lidocaine (1%) was injected under the scalp and a midline incision was made to expose the skull. To transduce ETs, an AAV (AAVretro-hSyn1-GCaMP6s-P2A-nls-tdTomato, titer: 5e^12^ gc/mL, Addgene), was injected into the right inferior colliculus of C57 mice through a small burr hole (5 mm posterior to bregma and 1.2 mm lateral). A glass micropipette containing the viral vector was lowered to depths of 0.3 and 0.8 mm using a micromanipulator (Kopf). A syringe pump (Nanoject III, Drummond) was used to inject 300 nL at each site at a rate of 10-15 nL/min. CTs were transduced with GCaMP6s through intracranial injection into the ACtx (AAV5-Syn-FLEX.GCaMP6s.WPRE.SV40, titer: 7e^12^ gc/mL, Addgene) or through transgenic crossing with the Ai148 mouse line. Post injection, the scalp was closed with nylon sutures and mice were administered with a subcutaneous injection of carprofen (5 mg/kg). Viruses were allowed to express for a minimum of three weeks prior to two-photon imaging. Cranial windows were created by stacking three round glass coverslips (one 4 mm, two 3 mm, #1 thickness, Warner Instruments), etched with piranha solution (sulfuric acid and hydrogen peroxide), then glued together with optically transparent, UV-cured adhesive (Norland Products, Warner Instruments). The skull was exposed by removing the dorsal portion of the scalp and periosteum and treated with etchant (C&B metabond) followed by 70% ethanol. A custom stainless steel headplate (iMaterialize, (Romero et al., 2020)) was then attached to the skull using dental cement (C&B metabond). A 3 mm circular craniotomy was made over right ACtx and the cranial window was placed in the craniotomy. An airtight seal was achieved using Kwik-Sil (World Precision Instruments) around the edge of the window prior to cementing into place. Vetbond (3M) was used to bind the surrounding tissue to the dental cement. Following the implant surgery, mice were administered with a subcutaneous injection of carprofen (5 mg/kg). Implants were allowed to heal for at least five days before the first day of imaging.

### Slice electrophysiology

Mice (P58-65) were first anesthetized with isoflurane and then immediately decapitated. Brains were rapidly removed and coronal slices (300 μm) containing the right ACtx were prepared in a cutting solution at 1 °C using a Vibratome (VT1200 S; Leica). The cutting solution, pH 7.4, ∼300 mOsm, contained the following (in mM): 2.5 KCl, 1.25 NaH_2_PO_4_, 25 NaHCO_3_, 0.5 CaCl_2_, 7 MgCl_2_, 7 Glucose, 205 sucrose, 1.3 ascorbic acid, and 3 sodium pyruvate (bubbled with 95% O_2_/5% CO_2_). The slices were immediately transferred and incubated at 34°C in a holding chamber for 40 min before recording. The holding chamber contained ACSF, containing the following (in mM): 125 NaCl, 2.5 KCl, 26.25 NaHCO_3_, 2 CaCl_2_, 1 MgCl_2_, 10 glucose, 1.3 ascorbic acid, and 3 sodium pyruvate, pH 7.4, ∼300 mOsm (bubbled with 95% O_2_/5% CO_2_). Post incubation, the slices were stored at room temperature until the time of recording. Whole-cell recordings in voltage- and current-clamp modes were performed on slices bathed in 31 °C carbogenated ACSF, which was identical to the incubating solution. Borosilicate pipettes (World Precision Instruments) were pulled into patch electrodes with 3–6 MΩ resistance (Sutter Instruments) and filled with a potassium-based intracellular solution, which was used for whole-cell current-clamp and contained (in mM): 128 K-gluconate, 10 HEPES, 4 MgCl_2_, 4 Na_2_ATP, 0.3 Tris-GTP, 10 Tris phosphocreatine, 1 EGTA, and 3 sodium ascorbate (pH = 7.25, 295 mOsm). For recording thalamically evoked Excitatory Postsynaptic Currents (EPSCs), a cesium-based internal solution containing (in mM): 126 CsCH_3_O_3_S, 4 MgCl_2_ 10 HEPES, 4 Na_2_ATP, 0.3 TrisGTP, 10 Tris-phosphocreatine, 1 CsEGTA, 1 QX-314, and 3 sodium ascorbate (pH = 7.25, 295 mOsm) was used. For electrophysiological recordings, we used a MultiClamp-700B amplifier equipped with Digidata-1440A A/D converter and Clampex (Molecular Devices). Data were sampled at 10 kHz and filtered at 4 kHz. Pipette capacitance was compensated and series resistance for recordings was lower than 25 MΩ. Series resistance (R_series_) was determined by giving a -5 mV voltage step for 50 ms in voltage-clamp mode (command potential set at -70 mV) and was monitored throughout the experiments. Recordings were excluded from further analysis if the series resistance changed by more than 20% compared to the baseline period. R_series_ was calculated by dividing the -5 mV voltage step by the peak current value generated immediately after the step in the command potential. Input resistance (R_input_) was calculated by giving a -5 mV step in voltage-clamp mode (command potential set at -70 mV), which resulted in transient current responses. The difference between baseline and steady-state hyperpolarized current (ΔI) was used to calculate R_input_ using the following formula: R_input_ = -5 mV/ΔI - R_series_. The average resting membrane potential (V_Rest_) was calculated by holding the neuron in voltage-follower mode (current clamp, at I = 0) immediately after breaking in and averaging the membrane potential over the next 20 sec. In experiments used to study intrinsic properties of neurons, we added the following drugs: 20 μΜ DNQX (AMPA receptor antagonist), 50 μΜ APV (NMDA receptor antagonist), and 20 μΜ SR 95531 Hydrobromide (Gabazine - a GABA_A_ receptor antagonist). Depolarizing current pulses (25 pA increments of 1-sec duration) in current clamp were injected to examine each neuron’s basic suprathreshold electrophysiological properties (baseline V_m_ was maintained at -70 mV, by injecting the required current, if necessary). Action potential (AP) width was calculated as the full width at the half-maximum amplitude of the first resulting action potential (AP) at rheobase. The AP threshold was measured in phase plane as the membrane potential at which the depolarization slope exhibited the first abrupt change (Δslope > 10 V/sec).

Light-evoked EPSCs and strontium quantal events (Sr^2+^-mEPSCs) were evoked by optogenetic stimulation of presynaptic axons. More specifically, a collimated blue LED light source (470 nm, Thorlabs) was directed through a diaphragm and a 40x microscope objective lens and restricted to a small spot in each stimulated cortical layer. Dual recordings targeted vertically aligned CT and ET pairs. The light intensity that elicited a reliable stable plateau response, maximal photostimulation, was determined on a cell pair basis with a short (0.15 ms duration) light pulse. EPSCs were recorded in voltage clamp mode at -70 mV (peak values were averaged over a 1 ms time window) in the presence of sodium channel blocker tetrodotoxin (TTX, 1μM) to isolate monosynaptic inputs and the potassium channel blocker 4-aminopyridine to depolarize the terminals (4-AP, 100 μM). Representative traces for figures were acquired by averaging 10 consecutive EPSCs. Sr^2+^-mEPSCs were recorded using a modified Sr^2+^-ACSF solution, which contained in mM: 125 NaCl, 2.5 KCl, 26 NaHCO_3_, 4 SrCl_2_, 4 MgCl_2_, 15 glucose, 1.3 ascorbic acid, and 3 sodium pyruvate, pH 7.4, ∼300 mOsm, oxygenated w/ 95% O_2_-5% CO_2_. Slices were incubated in this solution for 30 min prior to recording. All the other conditions were kept the same as described earlier. The same cutting solution, Cs^+^-based intracellular solution, and light activation were used. Clampfit (11.2) was used for Sr^2+^-mEPSCs analysis. A 400 ms window before LED stimulation (pre-LED) and another 400 ms window starting 100 ms after LED onset (post-LED) were analyzed. The amplitude of LED-evoked quantal events was calculated using the following equation 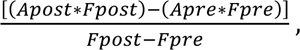 where *Apost* is the average amplitude of post-LED Sr^2+^-mEPSCs, *Fpost* is the average frequency of post-LED Sr^2+^-mEPSCs, *Apre* is the average amplitude of pre-LED events, and *Fpre* is the average frequency of pre-LED events (Whitt et al., 2022). Event detection was optimized by pre-processing the traces with a lowpass Gaussian filter (1kHz). Events (Sr^2+^-mEPSCs) with rise time < 3 ms and amplitude > 3 root mean square (RMS) were included. Cells with RMS noise > 2 or *Fpost* - *Fpre* < 2 Hz were excluded in the analysis.

### Noise and sham exposure

23-29 days after viral injections, mice (P50-55) were anesthetized with isoflurane (induction: 3% in oxygen, maintenance: 1.5-2% in oxygen) during a unilateral exposure via insertion of a plastic pipette tip into the left ear. The pipette tip was connected to a calibrated speaker (FT17H, Fostex, Middleton, WI). Core body temperature was maintained at ∼37°C with a heating pad and eyes were protected with ophthalmic ointment. Sham- and noise-exposed mice were treated in the same manner, but noise-exposed mice received 8-16 kHz octave broadband noise at 116 dB SPL for one hour. A DS360 Ultra-Low Distortion Function Generator (Stanford Research Systems-SRS) was used to generate bandwidth limiting using a 3-pole Butterworth filter. The speaker (Type 4231, Bruel & Kjaer) was calibrated with a 1/4-inch microphone (4954-B, Bruel & Kjaer) to accurately determine the voltage needed to generate the desired output.

### Auditory brainstem responses (ABRs)

ABR measurements were made from the left (exposed) ear and conducted in a sound-attenuating chamber (ENV-022SD; Med Associates). Subdermal electrodes were placed at the vertex (active), ventral to the right pinna (ground), and ventral to the left pinna (reference), with a calibrated speaker (CF-1; Tucker Davis Technologies) delivering acoustic stimuli to the left (exposed) ear through a pipette tip fixed to a plastic tube connected to the speaker. ABR measurements were obtained for tone bursts of 10, 16, 24, and 32 kHz in descending order from 80-0 dB SPL (or until 10 dB SPL below threshold). Threshold was defined as the lowest stimuli intensity that produced a Wave 1 response. A Wave 1 response waveform was identified and distinguished from non-response (noise) as the first consistent wave generated with decreased amplitude and increased latency as intensity decreased. ABR measurements were collected using BioSigRX software in response to 1 ms sound stimuli at a rate of 18.56 / sec, averaged over 512 repetitions and filtered using a 300-3,000 Hz bandpass filter.

### Two-photon imaging

At least three days prior to imaging, mice (P55-104) were acclimated to head fixation in the two-photon rig for 30 minutes a day. Imaging was carried out at three timepoints: one day pre-exposure (Day -1), one day post-exposure (Day 1), and seven days post-exposure (Day 7). ABRs were performed before each imaging session to measure peripheral sound responses. Mice with ABR click thresholds of 50 dB SPL or greater on Day -1 were excluded from the study. Noise-exposed mice were excluded if their click thresholds did not increase by at least 10 dB SPL from Day -1 to Day 1. Sham-exposed mice were excluded if their click thresholds exhibited an increase greater than 10 dB SPL from Day -1 to Day 1. On Day -1, widefield epifluorescence imaging was used to obtain a tonotopic map of ACtx to identify the location of primary area A1, as described previously (Romero et al., 2020). The two-photon field of view was then centered over A1 for all subsequent recordings. Two-photon excitation was delivered by an Insight X3 laser tuned to 940 nm (Spectra-Physics). A 16x/0.8NA water-immersion objective (Nikon) was used to obtain a 512 x 512 pixel field of view at 30 Hz with a Bergamo II Galvo-Resonant 8 kHz scanning microscope (Thorlabs). Scanning software (Thorlabs) and stimulus generation hardware (National Instruments) were synchronized with digital pulse trains. For both widefield and two-photon imaging, the microscope was rotated 40-55 degrees from the vertical axis to image the lateral portion of mouse ACtx while the mouse remained in an upright head position. Imaging was conducted in a dark, sound attenuating chamber and mice were monitored during experiments using infrared cameras (Genie Nano, Teledyne).

ETs were imaged 420-580 μm below the pial surface with a power of 45-130 mW. CTs were imaged 600-790 μm below the pial surface with a power of 111-150 mW. Fluorescence images were captured at either 1X (750 x 750 μm) or 2X (375 x 375 μm) digital zoom. Raw calcium videos were processed using Suite2P to accomplish motion correction, ROI detection, and spike deconvolution (Pachitariu M, 2016). All ROIs were visually validated, and any low quality or non-somatic ROIs were excluded. The same field of view was imaged at all three timepoints. Individual neurons were matched across days using ROIMatchPub, an open-source Matlab plugin (https://github.com/ransona/ROIMatchPub).

Each day of imaging included presentation of two stimulus sets. Eight pure tones (4 - 45 kHz in half-octave steps) were presented at four different intensities (20 - 80 dB SPL in 20 dB SPL steps). White noise bursts were presented at seven different intensities (20 - 80 dB SPL in 10 dB SPL steps). All stimuli were 50 ms in duration followed by a 2 s inter-trial interval. Stimuli were presented in pseudo-random order, and each stimulus was repeated 20 times.

### Two-photon imaging analysis

Deconvolved calcium traces were normalized for each cell by z-scoring the trace relative to the average baseline activity (500 ms prior to sound onset). Evoked stimulus responses were quantified as the mean z-scored activity 0 - 500 ms following sound onset. To determine neural responsiveness for each stimulus, z-scored activity in this stimulus response period was compared to 1000 samples of randomly shifted responses (5 - 20 second shifts in the trace). A cell was deemed significantly responsive to a stimulus if the evoked stimulus response was greater than the 98th percentile of the randomly shifted dataset (adapted from Khoury et al 2023). The best frequency of each cell was determined as the frequency that elicited the largest z-scored response after averaging across intensity. Intensity thresholds were determined as the lowest intensity that evoked a significant z-scored response.

### Stimulus decoding

We trained a multinomial logistic regression classifier to decode stimulus identity (either pure tone frequency at 80 dB SPL or white noise intensity) on a trial-by-trial basis. All neurons were pooled across mice for each exposure and day condition. Using 5-fold cross-validation, we trained the classifier with the mean evoked stimulus activity (0 - 500 ms following sound onset) of 200 randomly selected neurons. Decoding accuracy was quantified as the fraction of trials with correct stimulus classification in the held-out sample. Since our full dataset contained anywhere from 562 to 2687 neurons per group, we repeated this decoding strategy 2000 times, resampling (with replacement) a new population of 200 neurons on every iteration, to generated a resampled distribution of decoder accuracies for the population of neurons while ensuring decoding accuracies were not biased by the number of neurons in each group. We then calculated the mean and standard deviation of the decoder accuracy distribution in each exposure/day condition. P values were computed explicitly from the bootstrapped distributions of decoding accuracies (Day and Delgutte, 2016).

### Image processing

Image captioning in Figure 1C was achieved with Qcapture software. All images were imported and processed using ImageJ (https://imagej.nih.gov/ij/).

### Statistics

For statistical comparisons between two independent groups that passed the Shapiro-Wilk normality test, we used unpaired t-tests. Mann-Whitney Rank-sum tests were used for non-normally distributed data. For comparisons between multiple groups, we used N-way ANOVAs, corrected for multiple comparisons. Significance levels are denoted as *P < 0.05, **P < 0.01, ***P < 0.001, ****P < 0.0001. Group data are presented as mean ± SEM unless otherwise specified. See tables for detailed values and statistical tests (Tables 1-2).

**Table 2.** Statistical values for Supplemental Figures S1-S2.

| Figure | Statistical test | Main effect 1 | Main effect 2 | Main effect 3 | Interaction effect | N (mice) | Multiple comparison |
| --- | --- | --- | --- | --- | --- | --- | --- |
| S1A clicks | Mann-Whitney test |  |  | n/a |  |  | p < 0.0001 |
| S1A frequencies | 2-way ANOVA<br>Bonferroni | Exposure:<br>F (1,47) = 235.5<br>p < 0.0001 | Frequency:<br>F (2.598, 122.1) = 2.156<br>p = 0.1055 | n/a | F (3,141) = 3.93<br>p = 0.0099 | 1d SE = 24<br>1d NE = 25 | p < 0.0001 for all groups |
| S1B1 clicks | Mann-Whitney test |  |  | n/a |  |  | p < 0.0001 |
| S1B1 frequencies | 2-way ANOVA<br>Bonferroni | Exposure:<br>F (1,48) = 388.5<br>p < 0.0001 | Frequency:<br>F (2.855, 137.1) = 27.68<br>p < 0.0001 | n/a | F (3,144) = 3.839<br>p = 0.0111 | -1d NE = 25<br>1d NE = 25 | p < 0.0001 for all groups |
| S1B2 clicks | Mann-Whitney test |  |  | n/a |  |  | p = 0.5634 |
| S1B2 | 2-way ANOVA<br>Bonferroni | Exposure:<br>F (1,46) = 0.264<br>p = 0.6099 | Frequency:<br>F (2.778, 127.8) = 25.46<br>p < 0.0001 | n/a | F (3,138) = 0.8952<br>p = 0.4454 | -1d SE = 24<br>1d SE = 24 | ns for all groups |
| S1C1 | 3-way ANOVA | Exposure:<br>F (1,52) = 20.93<br>p < 0.0001 | Day:<br>F (1,52) = 11.06<br>p = 0.0016 | Intensity:<br>F (2.25,117) = 393.2<br>p < 0.0001 | F (7,364) = 8.586<br>p < 0.0001 | -1d SE = 14<br>1d SE = 14<br>-1d NE = 14<br>1d NE = 14 | n/a |
| S1C2 | 3-way ANOVA | Exposure:<br>F (1,52) = 29.84<br>p < 0.0001 | Day:<br>F (1,52) = 16.36<br>p = 0.0002 | Intensity:<br>F (3.153,163.9) = 314.5<br>p < 0.0001 | F (5,260) = 25.95<br>p < 0.0001 | -1d SE = 14<br>1d SE = 14<br>-1d NE = 14<br>1d NE = 14 | n/a |
| S1C3 | 3-way ANOVA | Exposure:<br>F (1,52) = 31.2<br>p < 0.0001 | Day:<br>F (1,52) = 17.9<br>p < 0.0001 | Intensity:<br>F (2.026,105.3) = 253.9<br>p < 0.0001 | F (7,364) = 8.467<br>p < 0.0001 | -1d SE = 14<br>1d SE = 14<br>-1d NE = 14<br>1d NE = 14 | n/a |
| S1C4 | 3-way ANOVA | Exposure:<br>F (1,52) = 20.63<br>p < 0.0001 | Day:<br>F (1,52) = 16.83<br>p = 0.0001 | Intensity:<br>F (1.636,85.08) = 205<br>p < 0.0001 | F (7,364) = 7.612<br>p < 0.0001 | -1d SE = 14<br>1d SE = 14<br>-1d NE = 14<br>1d NE = 14 | n/a |
| S1C5 | 3-way ANOVA | Exposure:<br>F (1,52) = 28.62<br>p < 0.0001 | Day:<br>F (1,52) = 22.12<br>p < 0.0001 | Intensity:<br>F (2.136,111.1) = 253.8<br>p < 0.0001 | F (6,132) = 11.39<br>p < 0.0001 | -1d SE = 14<br>1d SE = 14<br>-1d NE = 14<br>1d NE = 14 | n/a |
| S2A clicks | Mann-Whitney test |  |  | n/a |  |  | p = 0.0025 |
| S2A frequency | 2-way ANOVA<br>Bonferroni | Exposure:<br>F (1,23) = 32.09<br>p < 0.0001 | Frequency:<br>F (2.194, 50.46) = 3.018<br>p = 0.0532 | n/a | F (3,69) = 3.018<br>p = 0.0356 | 7d SE = 12<br>7d NE = 13 | 10 kHz, p = 0.0003<br>16 kHz, p < 0.0001<br>24 kHz, p < 0.0001<br>32 kHz, p = 0.0444 |
| S2B1 clicks | Mann-Whitney test |  |  | n/a |  |  | p = 0.001 |
| S2B1 frequency | Mixed-effects<br>Bonferroni | Exposure:<br>F (1,24) = 47.89<br>p < 0.0001 | Frequency:<br>F (2.801, 65.36) = 3.873<br>p = 0.0148 | n/a | F (3,70) = 3.069<br>p = 0.0334 | -1d NE = 13<br>7d NE = 13 | 10 kHz, p < 0.0001<br>16 kHz, p < 0.0001<br>24 kHz, p < 0.0001<br>32 kHz, p = 0.0063 |
| S2B2 clicks | Mann-Whitney test |  |  | n/a |  |  | p = 0.3261 |
| S2B2 frequency | 2-way ANOVA<br>Bonferroni | Exposure:<br>F (1,22) = 0.0599<br>p = 0.8089 | Frequency:<br>F (2.435, 53.56) = 7.606<br>p = 0.0006 | n/a | F (3,66) = 0.3511<br>p = 0.7885 | -1d SE = 12<br>7d SE = 12 | ns for all groups |
| S2C1 | 3-way ANOVA | Exposure:<br>F (1,20) = 27.6<br>p < 0.0001 | Day:<br>F (1,20) = 4.352<br>p = 0.0016 | Intensity:<br>F (1.954,39.07) = 189<br>p < 0.0001 | F (7,140) = 2.734<br>p = 0.0109 | -1d SE = 6<br>7d SE = 6<br>-1d NE = 6<br>7d NE = 6 | n/a |
| S2C2 | 3-way ANOVA | Exposure:<br>F (1,20) = 16.63<br>p = 0.0006 | Day:<br>F (1,20) = 3.692<br>p = 0.069 | Intensity:<br>F (1.852,37.03) = 67.87<br>p < 0.0001 | F (5,100) = 1.543<br>p = 0.1833 | -1d SE = 6<br>7d SE = 6<br>-1d NE = 6<br>7d NE = 6 | n/a |
| S2C3 | 3-way ANOVA | Exposure:<br>F (1,20) = 19.92<br>p = 0.0002 | Day:<br>F (1,20) = 3.485<br>p = 0.0767 | Intensity:<br>F (2.058,41.15) = 79.58<br>p < 0.0001 | F (6,120) = 1.949<br>p = 0.0784 | -1d SE = 6<br>7d SE = 6<br>-1d NE = 6<br>7d NE = 6 | n/a |
| S2C4 | 3-way ANOVA | Exposure:<br>F (1,20) = 15.3<br>p = 0.0009 | Day:<br>F (1,20) = 3.958<br>p = 0.0605 | Intensity:<br>F (2.063,41.26) = 85.47<br>p < 0.0001 | F (7,140) = 4.634<br>p = 0.0001 | -1d SE = 6<br>7d SE = 6<br>-1d NE = 6<br>7d NE = 6 | n/a |
| S2C5 | 3-way ANOVA | Exposure:<br>F (1,18) = 13.38<br>p = 0.0018 | Day:<br>F (1,18) = 2.074<br>p = 0.167 | Intensity:<br>F (1.879,33.83) = 54.56<br>p < 0.0001 | F (6,108) = 3.482<br>p = 0.0035 | -1d SE = 6<br>7d SE = 6<br>-1d NE = 6<br>7d NE = 6 | n/a |

## Results

To induce a stereotyped hearing loss, we followed standard approaches (Methods) and exposed mice to 8-16 kHz octave broadband noise at 116 dB SPL for 1 hour (noise-exposed, NE). As a control, we sham-exposed (SE) an additional cohort of littermate mice that underwent identical procedures to NE but without the presence of noise. To assess hearing thresholds pre and post noise- or sham-exposure (1-day and 7-day), we measured auditory brainstem responses (ABRs) (**Figure S1A-C**), which reflect the synchronous activity of auditory nuclei in response to sound arriving from the auditory nerve (wave I) to the inferior colliculus (wave V) (**Figure S1D**). We found significantly elevated ABR thresholds and reduced wave I amplitudes in both one day (**Figure S1A-D**) and 7 days after NE (**Figure S2A-C**) across all frequencies. ABR thresholds and wave I amplitudes were not affected in SE mice (**Figure S1A-D**). Together, these data suggest that our noise-exposure protocol results in threshold shifts that persist up to at least one week post ΝΕ.

### Transient shift in TC synaptic strength from CT and ET equivalent to CT dominant one day after NE

Communication between the ACtx and the auditory thalamus, the medial geniculate body (MGB), is mediated by reciprocally connected CT and TC pathways. TC neurons in the MGB strongly innervate neurons in the ACtx, including ETs and CTs (Hooks et al., 2013; Yamawaki and Shepherd, 2015). Yet, how NIHL affects TC synaptic strength onto ETs and CTs remains unknown.

To address this question, we performed dual patch-clamp recordings by simultaneously patching ETs and CTs from the same brain slice, while photoactivating TC axons. This recording configuration allowed us to calculate the change in the ratio of the synaptic strength of TC→CT over TC→ET synapses (CT/ET) in the same brain slice under identical experimental conditions. Although it is not optimal to compare absolute EPSC amplitudes with optogenetically-induced EPSC amplitudes, because of variable infection efficiency per mouse/brain area, we can meaningfully apply this technique to detect noise-induced changes in the relative strength of TC→CT vs. TC→ET synapses.

To study TC→CT and TC→ET synapses, we drove expression of channelrhodopsin (ChR2) in CAMKII-expressing MGB neurons by injecting AAV-CaMKII-ChR2-YFP into the MGB (**Figure 1A-C**, Methods). To stimulate ChR2-expressing TC axons, we used a collimated blue LED light source, directed through a diaphragm and a 40x microscope objective lens. We recorded monosynaptic photo-evoked AMPA receptor (AMPAR) EPSCs simultaneously from vertically aligned ETs (green) and CTs (red) (**Figure 1 A-C**). To selectively label ETs (green) and identify the boundaries of the ACtx, we injected green retrograde microspheres in the right inferior colliculus (IC, **Figure 1A, and C middle**) (Joshi et al., 2015; Joshi et al., 2016). To selectively label CTs (red), we injected Ntsr1-Cre transgenic mice, which selectively express Cre recombinase in CTs, with the Cre-dependent AAV vector AAV-FLEX-tdTomato (**Figure 1C right**). All electrophysiological experiments were performed in the presence TTX to block spikes, and 4-AP to enhance presynaptic depolarization (see Methods). Because CT dendrites are primarily limited to the deep cortical layers, we restricted our comparisons to L5-6 photostimulation (**Figure 1B**).

One day after noise exposure, we found a dramatic shift of the TC input from equivalent (ratio ∼ 1) to CT dominant (ratio > 5) (two-way ANOVA, main effect for exposure, F = 5.44, p = 0.02; main effect for day, F = 2.25, p = 0.14). 7 days after NE, we observed a recovery of TC inputs to a CT/ET equivalent state (**Figure 1D,E3-E4**). Together, these results suggest that noise exposure leads to a dramatic change in auditory TC input favoring a greater thalamic excitation of CTs compared to ETs one day after NIHL, which recovers seven days after trauma.

### Transient increase in the *q* on TC→CT synapses 1 day after NE

To understand the underlying mechanisms of this shift towards greater relative thalamic excitation in TC→CT compared to TC→ET synapses, we next investigated the synaptic properties of these synapses and the intrinsic excitability of CTs and ETs after noise trauma. Because these experiments were performed in TTX and 4-AP, and the ChR2 was activated to directly depolarize terminals, the probability of release (*p*) is likely to be saturated at all synapses (*n*). We therefore chose to assess the quantal size (*q*), first, in TC→CT synapses. To evaluate the amplitude of quantal events, we replaced calcium (Ca^2+^) with strontium (Sr^2+^) in the bath solution (**Figure 2A-D**). Sr^2+^ desynchronizes the evoked neurotransmitter release, thus allowing the analysis of quantal events from the stimulated synapses (**Figure 2A-D**) (Oliet et al., 1996; Kouvaros et al., 2020). In response to maximal photostimulation of L5-6, we found a significant increase in the average amplitude of quantal events from the stimulated TC→CT synapses one day after NE (unpaired t-test, p = 0.04; **Figure 2A-B**), which recovered 7 days after NE (unpaired t-test, p = 0.68; **Figure 2C-D**). This increase in *q* likely reflects an increase in TC→CT synaptic strength due to either postsynaptic changes or changes in vesicle content. Consistent with these findings, we found that the average L5-6 TC→CT EPSCs (average EPSCs elicited after L5-6 maximal photostimulation) were significantly enhanced 1 day after NE compared to SE (Mann-Whitney test, p = 0.004; **Figure 2E,F**), which then recovered 7 days after exposure (unpaired t-test, p = 0.94; **Figure 2G,H**). Together, these results suggest that the increase in *q* in TC→CT synapses after noise trauma contributes to the greater relative thalamic excitation in TC→CT compared to TC→ET after noise trauma.

**Figure 2.**
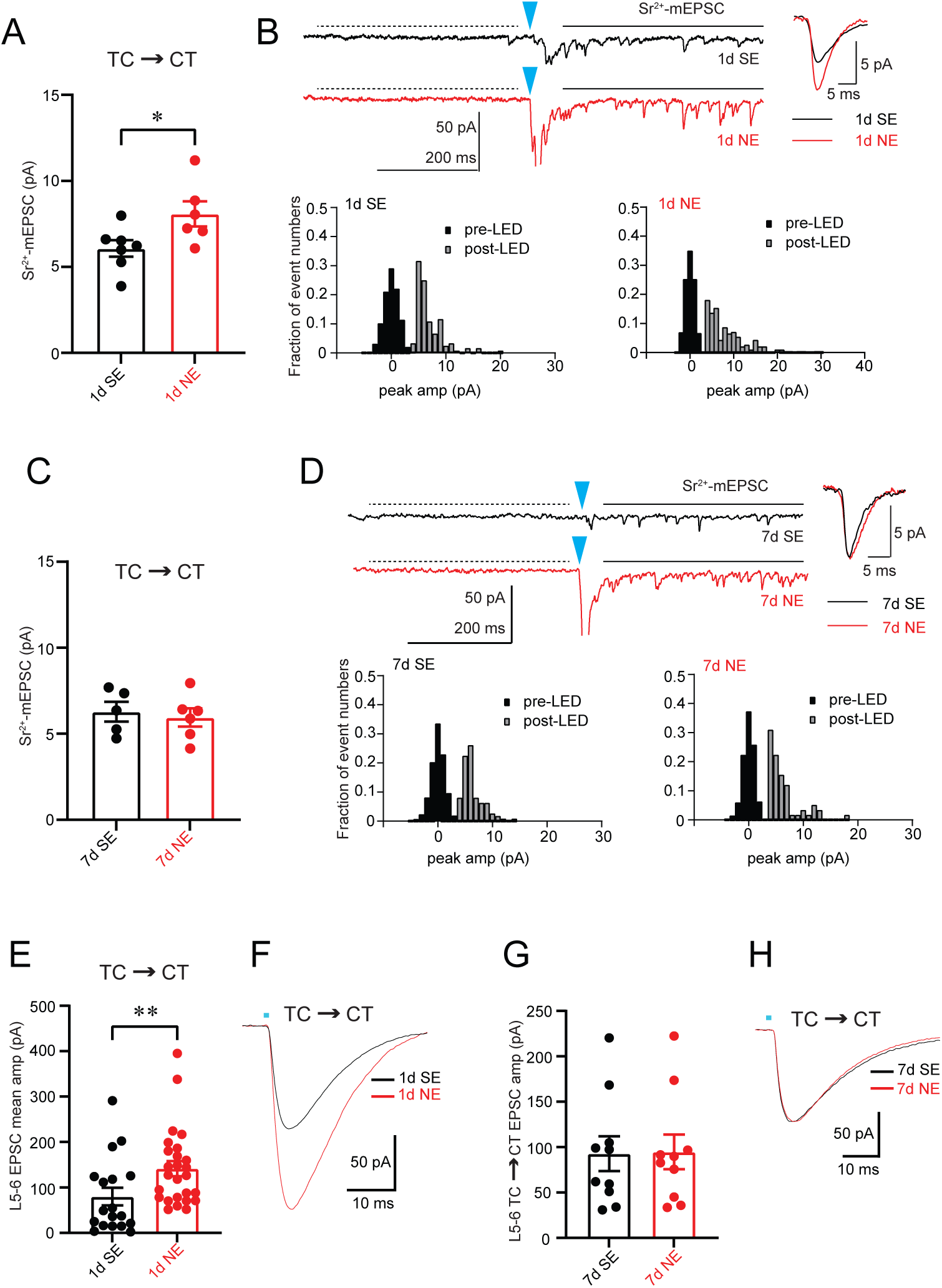
Transient increase in the *q* of TC→CT synapses 1 day after NE. **(A, C)** Average amplitude of light-evoked quantal EPSCs (Sr^2+^-mEPSCs) in CTs in response to L5-6 thalamocortical maximal photostimulation in 1d (A) and 7d (C), post SE (black) and NE (red) mice (1d SE: 7 cells/3 mice; 1d NE: 6 cells/3 mice; 7d SE: 5 cells/2 mice; 7d NE: 6 cells/2 mice). Asterisks indicate significant differences (*p<0.05, unpaired t-test). **(B, D)** Representative Sr^2+^-mEPSCs traces (top left) and representative average traces of quantal Sr^2+^-mEPSCs traces (top right). Amplitude histogram of events before (background noise) and after stimulation from the same cell (bottom) in 1d (B) and 7d (D) after SE (black) and NE (red) mice. The arrowhead indicates the onset of light stimulus. The dotted line indicates 400ms time window before stimulus (pre-LED). The solid line represents a 400ms time window, which started 100 ms after the stimulus (post-LED) and was used to analyze the amplitude of Sr^2+^-mEPSCs (1d SE: 7 cells/3 mice; 1d NE: 6 cells/3 mice; 7d SE: 5 cells/2 mice; 7d NE: 6 cells/2 mice). **(E, G)** Summary graph of the average CT EPSC amplitudes in response to L5-6 thalamocortical maximal photostimulation in 1d (D) and 7d (F) SE (black) and NE (red). (1d SE: 18 cells/6 mice; 1d NE: 26 cells/9 mice; 7d SE: 10 cells/6 mice; 7d NE: 10 cells/6 mice). Asterisks indicate significant differences (**p<0.01, Mann-Whitney test). **(F, H)** Representative traces of L6 CT EPSCs in response to L5-6 thalamocortical maximal photostimulation in 1d SE (E, black line) and 1d NE (E, red line) and 7d SE (G, black line) and 7d NE (G, red line). Detailed statistical values are listed in Table 1.

### Transient increase in CT suprathreshold and transient increase in subthreshold intrinsic excitability 1 day after NE

Next, we investigated the effect of noise trauma on CT intrinsic excitability. Neither input resistance (R_input_) nor action potential width or threshold (AP_width_, AP_thres_) were affected either at 1 or 7 days after NE (**Figure 3A,C-D**). We found that the resting membrane potential (V_rest_) was more hyperpolarized 1 day after NE (two-way ANOVA, main effect for exposure, F = 19.81, p < 0.0001; main effect for day, F = 9.40, p = 0.003; exposure x day interaction, F = 6.91, p = 0.01; **Figure 3B**). However, we found that the overall firing frequency in response to depolarization was significantly increased 1 day after NE (two-way ANOVA, main effect for exposure, F = 9.45, p = 0.004; exposure x current steps interaction, F = 3.82, p < 0.0001; **Figure 3E-F**), suggesting an increased suprathreshold excitability of CTs concurrent with the increased TC→CT input. There were no differences in CT intrinsic excitability between SE and NE mice 7 days after exposure (two-way ANOVA, main effect for exposure, F = 2.64, p = 0.11; exposure x current steps interaction, F = 0.91, p = 0.52; **Figures 3G-H**). Taken together, these results suggest that although there is a hyperpolarization on V_rest_ after NE, a combination of intrinsic and synaptic mechanisms that increase TC→CT synaptic strength and CT suprathreshold excitability likely contribute to the shift of TC input from equivalent between CT and ET to CT dominant one day after noise exposure.

**Figure 3.**
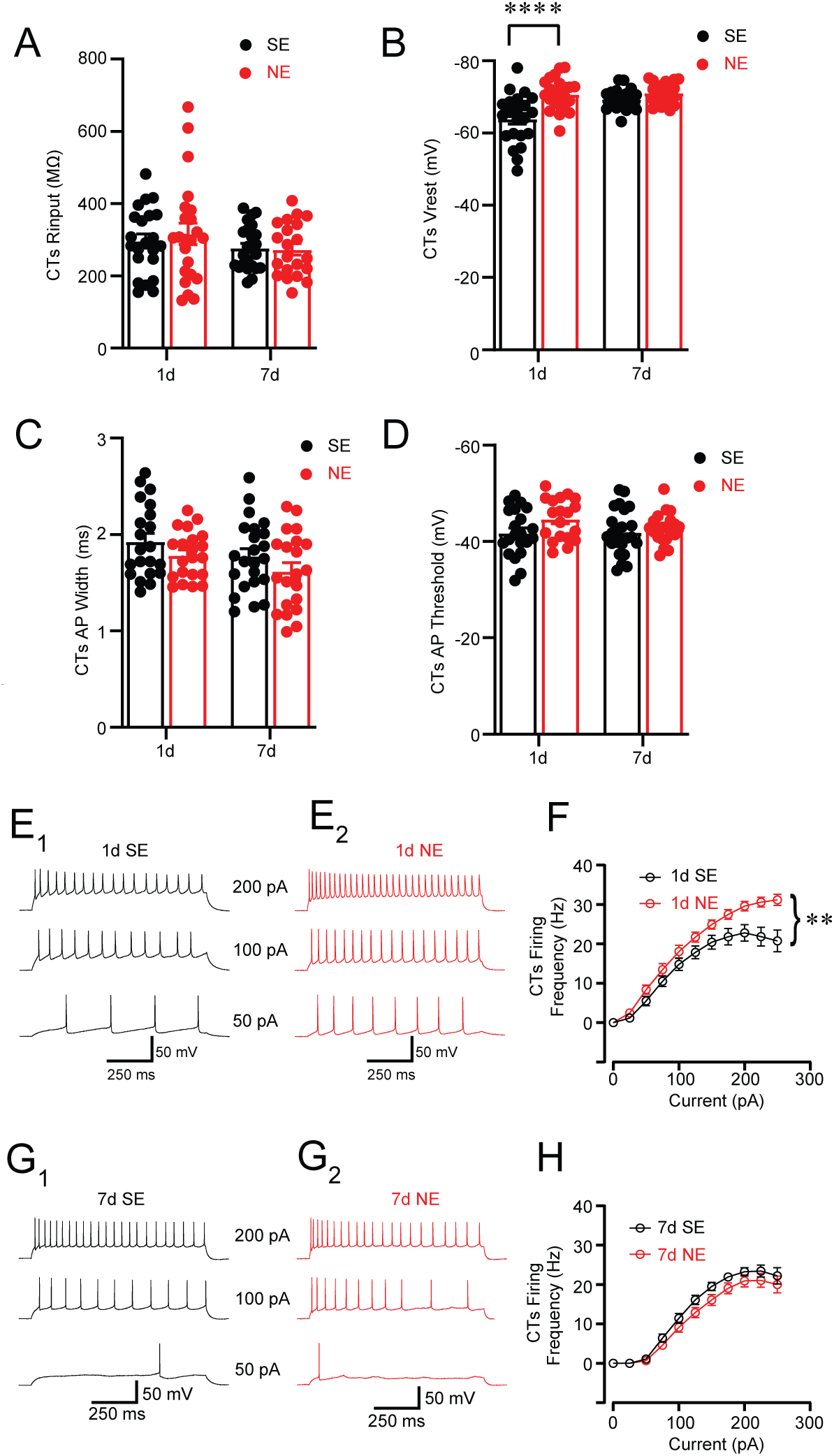
Transient increase in suprathreshold CT intrinsic excitability 1 day after NE. **(A)** Average CT input resistance (R_input_) 1d and 7d after SE (black) or NE (red) (1d SE: 22 cells/8 mice; 1d NE: 23 cells/5 mice; 7d SE: 23 cells/6 mice; 7d NE: 21 cells/4 mice). **(B)** Average CT resting membrane potential (V_rest_) 1d and 7d after SE (black) or NE (red) (1d SE: 22 cells/8 mice; 1d NE: 23 cells/5 mice; 7d SE: 23 cells/6 mice; 7d NE: 21 cells/4 mice). **(C)** Average CT AP width 1d and 7d after SE (black) or NE (red) (1d SE: 21 cells/8 mice; 1d NE: 21 cells/5 mice; 7d SE: 23 cells/6 mice; 7d NE: 21 cells/4 mice). **(D)** Average CT AP threshold of 1d and 7d after SE (black) or NE (red) (1d SE: 21 cells/8 mice; 1d NE: 19 cells/5 mice; 7d SE: 23 cells/6 mice; 7d NE: 21 cells/4 mice). **(E_1_, E_2_, G_1_, G_2_)** Representative traces of CT firing in response to depolarizing current (50, 100, 200 pA current injections), 1d SE (E_1_) vs. 1d NE (E_2_); and 7d SE (G_1_) vs. 7d NE (G_2_). **(F, H)** Average firing frequency as a function of injected current amplitude, 1d SE vs. 1d NE (F), and 7d SE vs. 7d NE (H). Current injections from 25 to 250 pA with an increment of 25 pA (1d SE: 21 cells/8 mice; 1d NE: 21 cells/5 mice; 7d SE: 23 cells/6 mice; 7d NE: 21 cells/4 mice). Asterisks indicate significant differences (**p<0.01, two-way ANOVA and Bonferroni correction for multiple comparisons). Detailed statistical values are listed in Table 1.

### No changes in the *q* of TC→ET synapses after NE

We next assessed TC→ET synapses using a similar electrophysiological approach as in **Figure 2**. We found no changes in *q* either at 1 day or 7 days after exposure (1 day, Mann-Whitney test, p = 0.64; 7 days, Mann-Whitney test, p > 0.9999; **Figure 4A-D**). Moreover, we found that the average TC→ET EPSCs were significantly reduced one day after NE compared to SE (two-way ANOVA, main effect for exposure, F = 8.32, p = 0.005; main effect for day, F = 0.05, p = 0.82; exposure x day interaction, F = 4.97, p = 0.03; **Figure 4E-F_1_**). This reduction was recovered 7 days after exposure (**Figure 4E,F_2_**). Given that *p* is likely saturated in TC→ET synapses, and *q* is unaltered, the reduction in L5-6 TC→ET EPSCs might be due to a decrease in *n*, or to the previously mentioned limitations of this experimental setup in assessing changes in synaptic strength, for changes in observed synaptic strength might be due to inconsistent degree of AAV infection across animals (Discussion).

**Figure 4.**
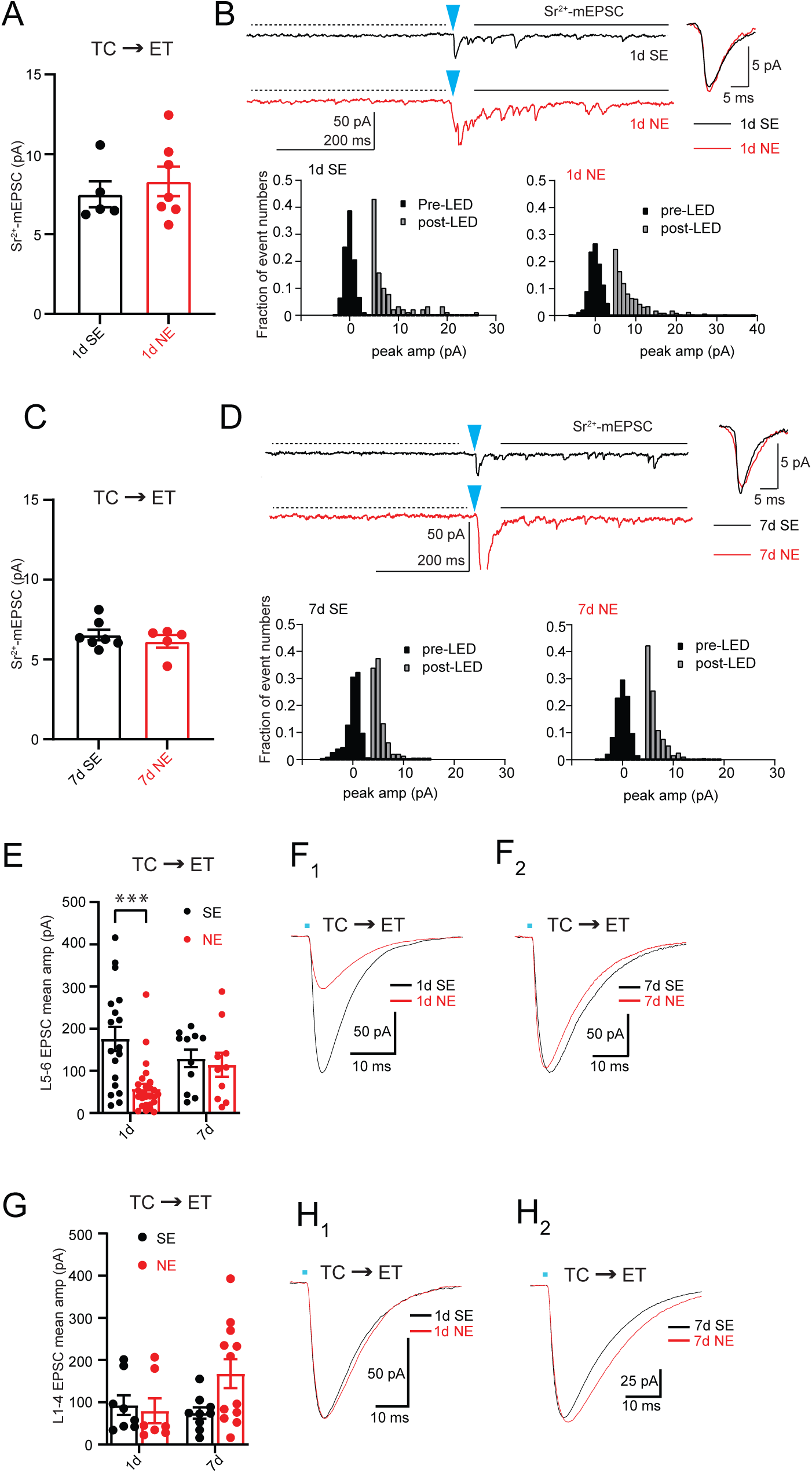
No changes in the *q* of TC→ET synapses after NE. **(A, C)** Average amplitude of Sr^2+^-mEPSCs in ETs in response to L5-6 thalamocortical maximal photostimulation in 1d (A) and 7d (B), post SE (black) and NE (red) (1d SE: 5 cells/4 mice; 1d NE: 7 cells/5 mice; 7d SE: 7 cells/2 mice; 7d NE: 5 cells/2 mice). **(B, D)** Representative Sr^2+^-mEPSCs traces (top left) and representative average traces of quantal Sr^2+^-mEPSCs traces (top right). Amplitude histogram of events before (background noise) and after stimulation from the same cell (bottom) in 1d (B) and 7d (D) after SE (black) and NE (red) mice. The arrowhead indicates the onset of light stimulus. The dotted line indicates 400ms time window before stimulus (pre-LED). The solid line represents a 400ms time window, which started 100 ms after the stimulus (post-LED) and was used to analyze the amplitude of Sr^2+^-mEPSCs (1d SE: 5 cells/4 mice; 1d NE: 7 cells/5 mice; 7d SE: 7 cells/2 mice; 7d NE: 5 cells/2 mice). **(E)** Representative traces of L5 ET EPSCs in response to L5-6 thalamocortical maximal photostimulation in in 1d SE (E_1_, black) vs. 1d NE (E_1_, red), 7d SE (E_2_, black) vs. 7d NE (E_2_, red) mice. **(F)** Summary graph of the average L5 ET EPSC amplitudes in response to L1-4 thalamocortical maximal photostimulation in 1d SE vs. 1d NE, or 7d SE vs. 7d NE mice (1d SE: 8 cells/3 mice; 1d NE: 7 cells/4 mice; 7d SE: 9 cells/5 mice; 7d NE: 12 cells/4 mice). **(G)** Representative traces of L5 ET EPSCs in response to L1-4 thalamocortical maximal photostimulation in 1d SE (G_1_, black) vs. 1d NE (G_1_, red), 7d SE (G_2_, black) vs 7d NE (G_2_, red) mice. Detailed statistical values are listed in Table 1.

Because ETs have extensive dendrites throughout the cortical column, and TC projections to ETs span several cortical layers (Harris and Shepherd, 2015; Ji et al., 2016), we also averaged TC→ET EPSCs elicited after photostimulating L1-4 (L1-4 TC EPSCs). No significant changes were found at either 1 day or 7 days after NE (two-way ANOVA, main effect for exposure, F = 1.88, p = 0.18; main effect for day, F = 1.43, p = 0.24; exposure x day interaction, F = 3.35, p = 0.08; **Figure 4G-H**). Together, these results suggest that there is a cell-type-specific increase in *q* in TC→CT synapses that likely contributes to the greater thalamic excitation of CTs compared to ETs one day after NIHL.

### No changes in ET intrinsic excitability after NE

After establishing that NIHL differentially affects TC onto ETs vs. CTs, we next investigated whether noise exposure also alters the intrinsic excitability of these neurons in a cell-type-specific manner. R_input_, V_rest_, AP_width_, AP_threshold_ and current-firing frequency function did not change either at 1 day or 7 days post NE in ETs (**Figure 5A-H**). This finding supports cell-type-specific intrinsic plasticity in deep layer neurons after NIHL, whereby ETs do not show plasticity while CTs show a transient decrease in subthreshold excitability and a transient increase in suprathreshold excitability. Together, these results suggest that the cell-type-specific increase in *q* in TC→CT synapses and in suprathreshold CT intrinsic excitability likely contribute to the greater thalamic excitation of CTs compared to ETs one day after NIHL.

**Figure 5.**
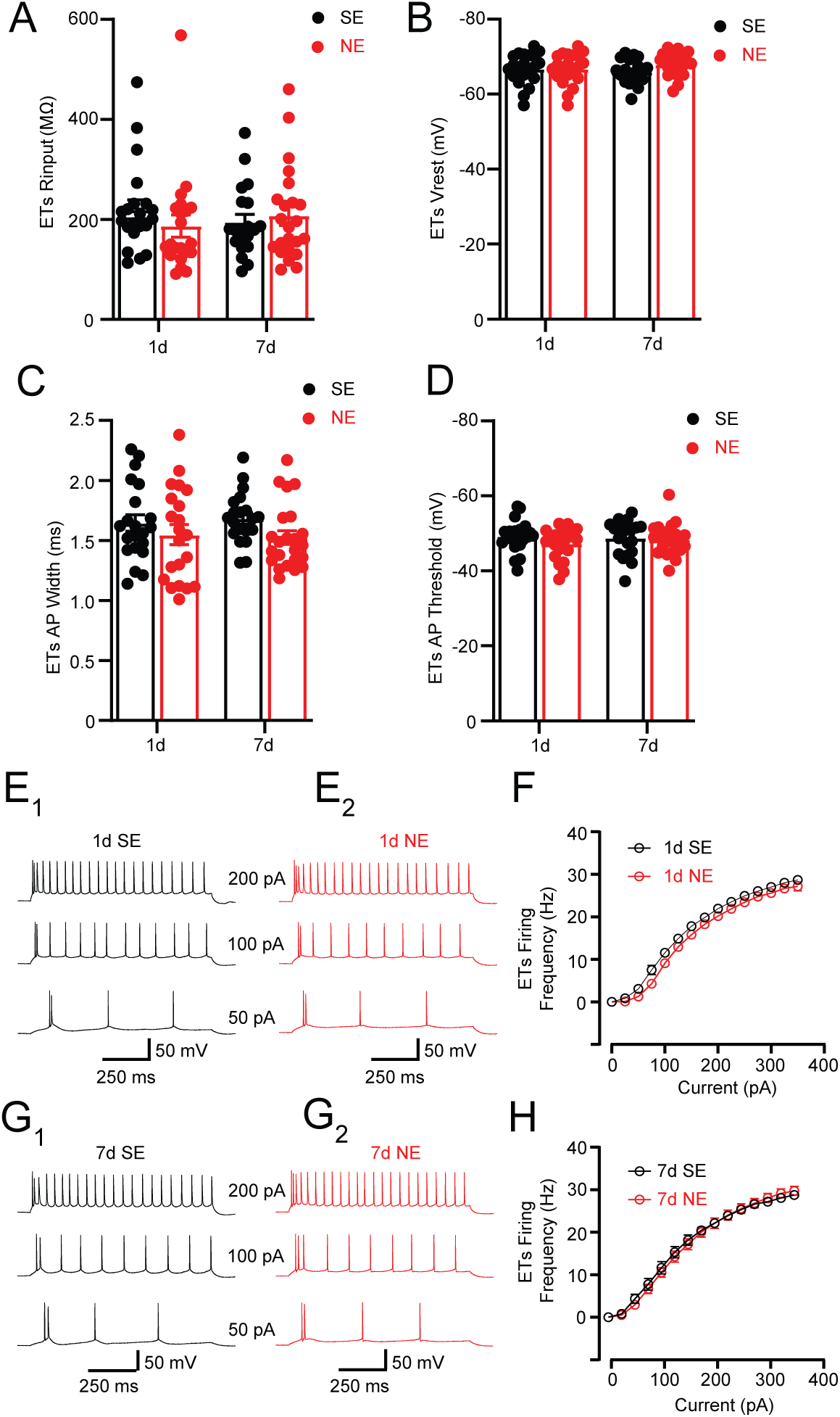
No changes in ET intrinsic excitability after NE. **(A)** Average ET R_input_ 1d and 7d after SE (black) or NE (red) (1d SE: 21 cells/6 mice; 1d NE: 21 cells/5 mice; 7d SE: 20 cells/5 mice; 7d NE: 23 cells/6 mice). **(B)** Average ET V_rest_ 1d and 7d after SE (black) or NE (red) (1d SE: 21 cells/6 mice; 1d NE: 21 cells/5 mice; 7d SE: 20 cells/5 mice; 7d NE: 23 cells/6 mice). **(C)** Average ET AP width 1d and 7d after SE (black) or NE (red) (1d SE: 21 cells/6 mice; 1d NE: 21 cells/5 mice; 7d SE: 20 cells/5 mice; 7d NE: 23 cells/6 mice). **(D)** Average ET AP threshold 1d and 7d after SE (black) or NE (red) (1d SE: 21 cells/6 mice; 1d NE: 21 cells/5 mice; 7d SE: 20 cells/5 mice; 7d NE: 23 cells/6 mice). **(E_1_, E_2_, G_1_, G_2_)** Representative traces of ET firing in response to depolarizing current injection (50, 100, 200 pA), 1d SE (E_1_) vs. 1d NE (E_2_); and 7d SE (G_1_) vs. 7d NE (G_2_). **(F, H)** Average firing frequency as a function of injected current amplitude, 1d SE vs. 1d NE (F), and 7d SE vs. 7d NE (H). Current injections from 25 to 350 pA with an increment of 25 pA. (1d SE: 21 cells/6 mice; 1d NE: 21 cells/5 mice; 7d SE: 20 cells/5 mice; 7d NE: 23 cells/6 mice). Detailed statistical values are listed in Table 1.

### Changes in CT sound response properties persist 7 days after NE

To examine the consequences of NIHL in ET and CT sound response properties, we recorded the sound-evoked activity of neural populations using two-photon (2P) calcium imaging. This technique allowed us to longitudinally image the same populations of genetically defined cell-types across time before and after noise exposure and characterize changes in sound response properties. Head-fixed, awake mice passively listened to auditory stimuli, and we recorded responses one day prior to exposure, then again one and seven days after exposure, matching our previous electrophysiological studies. We selectively expressed GCaMP6s in CTs to examine how CT responses to sounds change following NE (**Figure 6A**, see Methods). We imaged a total of 4943 CTs across 8 unique fields of view in SE mice (-1d: 1649 neurons; 1d: 1681 neurons; 7d: 1613 neurons) and 7731 CTs across 14 unique fields of view in NE mice (-1d: 2687 neurons; 1d: 2546 neurons; 7d: 2498 neurons). We then determined which neurons were significantly active during sound presentation and used these sound-responsive neurons for all subsequent analysis (see Methods).

**Figure 6.**
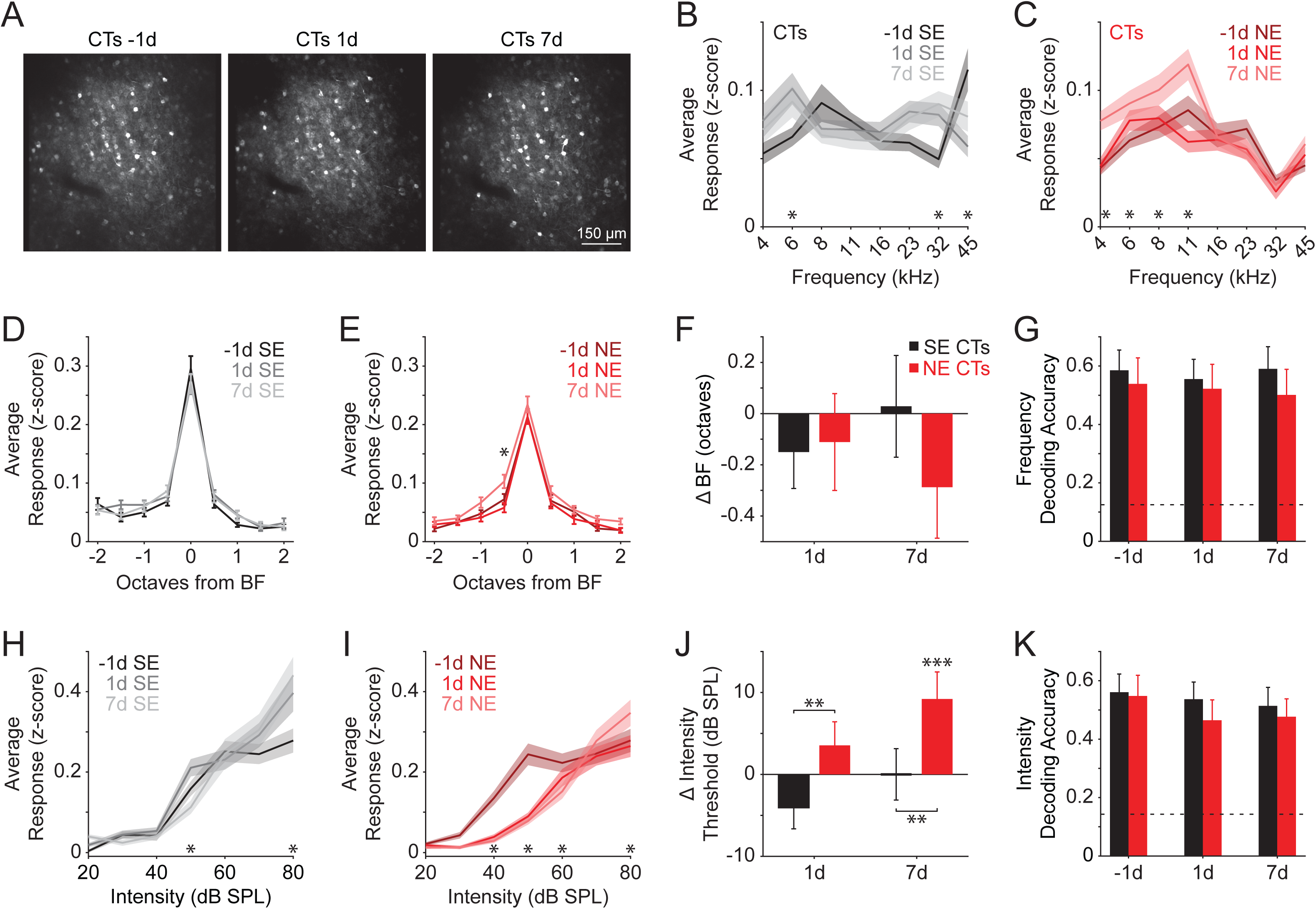
Changes in CT sound response responses persist 7 days after NE. **(A)** Example two-photon field of view from CTs imaged before and after exposure. **(B)** Average responses to pure tones across all CTs in SE mice (-1d: n=507 neurons; 1d: n=460 neurons; 7d: n=432 neurons). Asterisks indicate significant pairwise differences. **(C)** Same as (**A**) for NE mice (-1d: n=713 neurons; 1d: n=638 neurons; 7d: n=646 neurons). Asterisks indicate significant pairwise differences. **(D)** Tuning curves aligned to best frequency (BF) in SE mice. **(E)** Same as (**D**) for NE mice. Asterisks indicate significant pairwise differences (*p<0.05, Bonferroni corrected for multiple comparisons). **(F)** Change in BF of matched cells in SE (black) and NE (red) mice (1d SE: n=40 neurons; 1d NE: n=36 neurons; 7d SE: n=35 neurons; 7d NE: n=40 neurons). **(G)** Accuracy of a multinomial logistic regression classifier trained to decode the frequency of pure tones at 80 dB SPL. Dashed line represents chance level. Error bars represent standard deviation of decoding iterations. **(H)** Average responses to white noise bursts at different intensities across all ETs in SE mice (- 1d: n=438 neurons; 1d: n=460 neurons; 7d: n=411 neurons). Asterisks indicate significant pairwise differences (Bonferroni corrected for multiple comparisons). **(I)** Same as (**H**) in NE mice (-1d: n=636 neurons; 1d: n=559 neurons; 7d: n=587 neurons). Asterisks indicate significant pairwise differences (Bonferroni corrected for multiple comparisons). **(J)** Change in intensity thresholds across days in SE (black) and NE (red) mice (1d SE: n=27 neurons; 1d NE: n=31 neurons; 7d SE: n=29 neurons; 7d NE: n=25 neurons). Asterisks indicate significant pairwise differences (**p<0.01, ***p<0.001, Bonferroni corrected for multiple comparisons). **(K)** Accuracy of a multinomial logistic regression trained to decode sound intensity. Dashed line represents chance level. Error bars represent standard deviation of decoding iterations. Detailed statistical values are listed in Table 1.

We first examined frequency responses by presenting pure tones of varying frequencies (4-45 kHz in 0.5 octave steps) and intensities (20-80 dB SPL in 20 dB SPL steps). Averaging across intensity, we initially observed that CTs exhibited unstable frequency encoding, as evidenced by fluctuations in average pure tone responses in SE animals (two-way ANOVA, main effect for day, F = 0.66, p = 0.52; frequency × day interaction, F = 3.68, p < 0.0001; **Figure 6B**). Nevertheless, we observed strong changes in CT sound responses following NE (two-way ANOVA, main effect for day, F = 11.62, p < 0.0001; frequency × day interaction, F = 2.78, p = 0.0004; **Figure 6C**). CTs exhibited increased population responses to low frequency sounds (4-11 kHz) 7 days after NE. Next, we compared average population tuning curves by aligning activity to each neuron’s best frequency (BF) and comparing these aligned tuning curves across days. Frequency tuning curves remained stable in SE mice (two-way ANOVA, main effect for day, F = 0.52, p = 0.59; frequency × day interaction, F = 0.81, p = 0.67; **Figure 6D**). NE induced a slight but significant increase in responses 0.5 octaves below BF (two-way ANOVA, main effect for day, F = 9.36, p < 0.0001; frequency × day interaction, F = 0.52, p = 0.94; **Figure 6E**). To investigate if individual neurons altered their tuning preferences following noise exposure, we matched the same cells across days and quantified changes in BF compared to baseline on day -1. While not significant, CTs displayed a trend to decrease BF following NE (two-way ANOVA, main effect for exposure, F = 1.00, p = 0.32; main effect for day, F = 1.14, p = 0.32; frequency × day interaction, F = 1.33, p = 0.27; **Figure 6F**). To examine how these changes in frequency responses impacted population encoding, we used a multinomial logistic regression model to predict stimulus frequency from neural activity. Decoding frequency identity from CT activity did not reveal any effect of noise exposure on population-level encoding (**Figure 6G**). Together, these results suggest that noise exposure does not profoundly change CT frequency tuning.

We next examined intensity tuning by presenting broadband noise bursts between 20-80 dB SPL (in 10 dB SPL steps). As with pure tone responses, CTs showed subtle changes in broadband noise responses following SE (two-way ANOVA, main effect for day, F = 3.54, p = 0.03; frequency × day interaction, F = 2.51, p = 0.003; **Figure 6H**). In contrast, there was a marked decrease in intensity responses following NE (two-way ANOVA, main effect for day, F = 13.87, p < 0.0001; frequency × day interaction term, F = 5.45, p < 0.0001; **Figure 6I**). Specifically, CTs exhibited decreased activity in response to low intensity noise (< 60 dB SPL) following NE, which did not recover by day 7. There was also a slight but significant increase in responses to 80 dB SPL noise 7 days after NE. Consequently, NE led to a significant increase in intensity thresholds (two-way ANOVA, main effect for exposure, F = 14.75, p = 0.0002; main effect for day, F = 4.165, p = 0.02; frequency × day interaction, F = 4.94, p = 0.008; **Figure 6J**). Despite these changes, decoding analyses did not reveal any significant changes in population encoding (**Figure 6K**). Together, this suggests that noise exposure leads to decreased sensitivity in CTs to low intensity noise that does not recover within a week.

### Transient changes in ET sound response properties 1 day after NE

We next examined the activity of ETs through selective expression of GCaMP6s (**Figure 7A**, see Methods). We imaged a total of 2337 ETs across 9 unique fields of view in SE mice (-1d: 705 neurons; 1d: 820 neurons; 7d: 812 neurons) and 1703 ET neurons across 8 unique fields of view in NE mice (-1d: 562 neurons; 1d: 565 neurons; 7d: 576 neurons). As before, only neurons that were significantly active during sound presentation were used for subsequent analyses.

**Figure 7.**
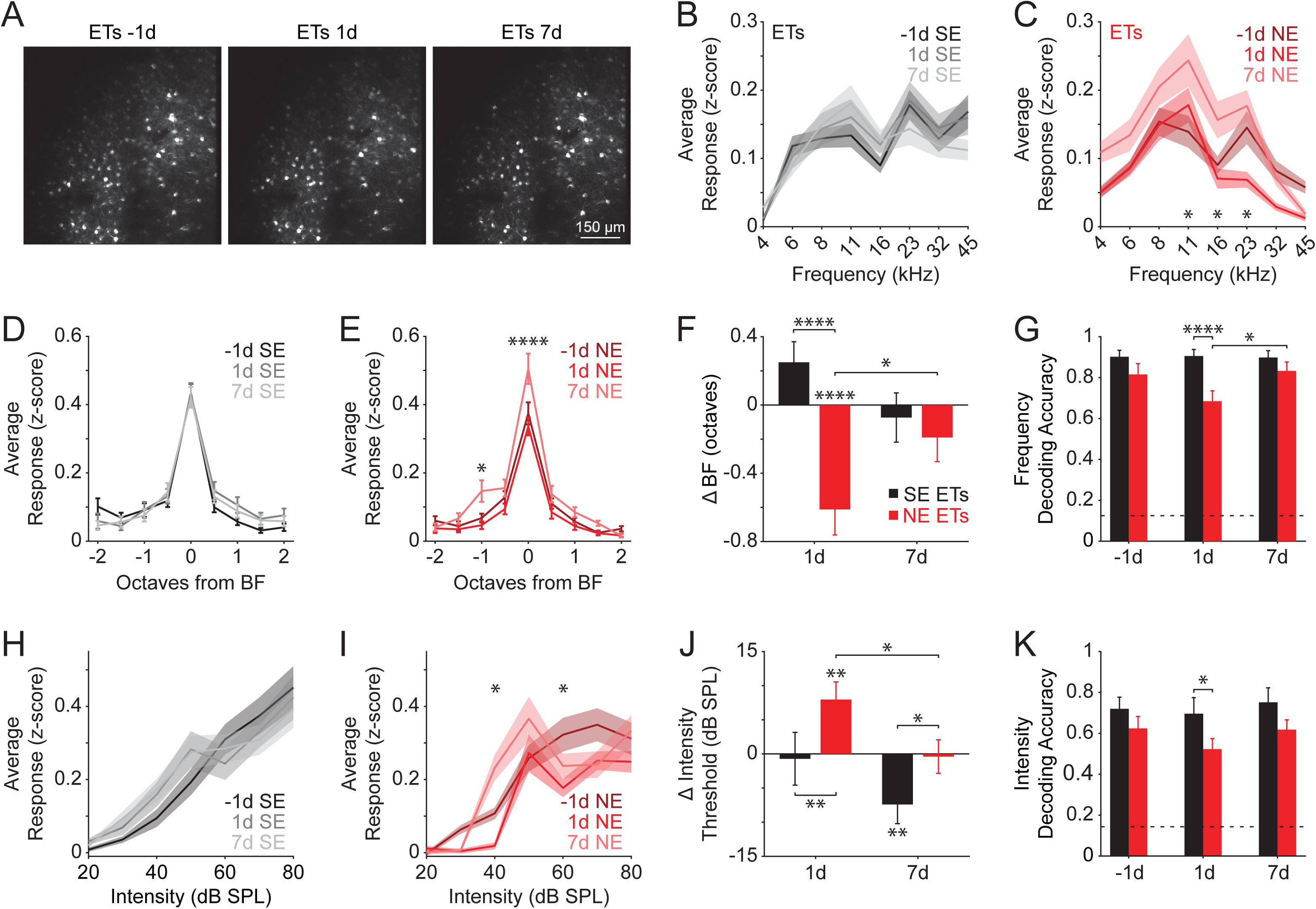
Transient shift in ET sound response properties one day after NE. **(A)** Example two-photon field of view from ETs imaged before and after exposure. **(B)** Average responses to pure tones across all ETs in SE mice (-1d: n=385 neurons; 1d: n=414 neurons; 7d: n=400 neurons). **(C)** Same as in (**A**) for NE mice (-1d: n=268 neurons; 1d: n=235 neurons; 7d: n=296 neurons). Asterisks indicate significant pairwise differences (Bonferroni corrected for multiple comparisons). **(D)** Tuning curves aligned to best frequency (BF) in SE mice. **(E)** Same as in (**D**) for NE mice. Asterisks indicate significant pairwise differences (*p<0.05, ****p<0.0001, Bonferroni corrected for multiple comparisons). **(F)** Change in BF of matched cells in SE (black) and NE (red) mice (1d SE: n=68 neurons; 1d NE: n=54 neurons; 7d SE: n=68 neurons; 7d NE: n=71 neurons). Asterisks indicate significant pairwise differences (*p<0.05, ****p<0.0001, Bonferroni corrected for multiple comparisons). **(G)** Accuracy of a multinomial logistic regression classifier trained to decode the frequency of pure tones at 80 dB SPL. Dashed line represents chance level. Error bars represent standard deviation of decoding iterations. Asterisks indicate significant pairwise differences (*p<0.05, ****p<0.0001, bootstrap test Bonferroni corrected for multiple comparisons). **(H)** Average responses to white noise bursts at different intensities across all ETs in SE mice (- 1d: n=331 neurons; 1d: n=270 neurons; 7d: n=299 neurons). **(I)** Same as (**H**) in NE mice (-1d: n=226 neurons; 1d: n=222 neurons; 7d: n=252 neurons). Asterisks indicate significant pairwise differences (Bonferroni corrected for multiple comparisons). **(J)** Change in intensity thresholds across days in SE (black) and NE (red) mice (1d SE: n=42 neurons; 1d NE: n=49 neurons; 7d SE: n=50 neurons; 7d NE: n=51 neurons). Asterisks indicate significant pairwise differences (*p<0.05, **p<0.01, Bonferroni corrected for multiple comparisons). **(K)** Accuracy of a multinomial logistic regression trained to decode sound intensity. Dashed line represents chance level. Error bars represent standard deviation of decoding iterations. Asterisks indicate significant pairwise differences (*p<0.05, bootstrap test Bonferroni corrected for multiple comparisons). Detailed statistical values are listed in Table 1.

We found that the average population responses to pure tones remained unchanged in SE mice (two-way analysis of variance (ANOVA), main effect for day, F = 0.68, p = 0.51; frequency × day interaction, F = 1.03, p = 0.41; **Figure 7B**). In contrast, NE led to a shift in the representation of frequencies across the ET population (two-way ANOVA, main effect for day, F = 19.35, p < 0.0001; frequency × day interaction, F = 1.93, p = 0.02; **Figure 7C**). One day after NE, the average response to 23 kHz pure tones significantly decreased compared to baseline. This reduction in activity then recovered to baseline levels by day 7 and was accompanied by a significant increase in response to lower frequency sounds (11-16 kHz). BF-aligned tuning curves were consistent across days in SE mice (two-way ANOVA, main effect for day, F = 0.58, p = 0.56; frequency × day interaction, F = 0.58, p = 0.90; **Figure 7D**). Conversely, NE led to significant changes in the average tuning curves (two-way ANOVA, main effect for day, F = 12.03, p < 0.0001; frequency × day interaction, F = 1.84, p = 0.02; **Figure 7E**). Specifically, 7 days after NE, ETs increased their responses at BF and one octave below BF (two-way ANOVA, main effect for exposure, F = 17.67, p < 0.0001; main effect for day, F = 2.23, p = 0.11; exposure × day interaction term, F = 11.34, p < 0.0001). This was accompanied by a significant decrease in BF on day 1 which then recovered by day 7 in NE animals (**Figure 7F**). Frequency decoding revealed that SE populations exhibited stable predictive accuracy across days, while NE led to a significant decrease in accuracy on day 1 and a subsequent recovery on day 7 (**Figure 7G**). These findings suggest that one day following NE, ETs reduce their activity at high frequencies and consequently shift to lower BFs, decreasing the ability to accurately decode stimuli from population activity. These perturbations then return to baseline seven days after NE.

We next examined whether ET intensity tuning changed following NE. As expected, noise responses were similar across days in SE mice (two-way ANOVA, main effect for day, F = 0.20, p = 0.82; intensity × day interaction, F = 0.94, p = 0.50; **Figure 7H**). We observed that NE led to significant changes in intensity responses (two-way ANOVA, main effect for day, F = 8.01, p = 0.0003; frequency × day interaction term, F = 2.90, p = 0.0005; **Figure 7I**). One day after NE, ETs displayed a loss of responses to low intensities (< 50 dB SPL) and a reduction in responses at higher intensities. On day 7, responses partially recovered but remained below baseline for 70 dB SPL. We then quantified changes in single neuron intensity thresholds with cells matched across imaging days (two-way ANOVA, main effect for exposure, F = 11.23, p = 0.0009; main effect for day, F = 6.56, p = 0.002; exposure × day interaction, F = 3.496, p = 0.03; **Figure 7J**). One day after NE, ETs displayed an increased threshold which reverted to baseline on day 7. We also observed a threshold decrease on day 7 in SE mice. Finally, we decoded sound intensity from population activity using multinomial logistic regression. Similar to our frequency decoding findings, we found that decoding accuracy transiently decreased on day 1 following NE (**Figure 7K**). Together, these findings suggest that ETs recover from transient changes in tuning to both frequency and intensity following noise exposure.

## Discussion

Our findings provide new insights into the cellular and circuit mechanisms underlying the cortical plasticity of deep-layer neurons in the auditory cortex (ACtx) following noise-induced hearing loss (NIHL). We identified distinct changes in the synaptic, intrinsic, and sound response properties of L5 ET and L6 CT neurons. These findings advance our understanding of ACtx plasticity after peripheral damage and suggest a transient change in the relative strength of thalamocortical inputs to ETs and CTs that may help maintain auditory function post-NIHL.

The observed changes in synaptic strength in thalamic input between ETs and CTs were assessed using optogenetic approaches, which can be influenced by variability in viral infection efficiency across animals and brain areas, and our recordings conditions where *p* is likely saturated (**Figures 2E-H and 4E-H**). However, simultaneous recordings from ETs and CTs, which mitigate these limitations, revealed a greater relative thalamic excitation of CTs compared to ETs one day after NIHL (**Figure 1**). This finding, along with the increased *q* in TC→CT synapses (**Figure 2A-D**) and the increased suprathreshold intrinsic CT excitability (**Figure 3E,F**), which are independent of the variability in viral infection and *p* saturation, support an increase in thalamocortical synaptic strength to CTs after noise trauma.

CTs form reciprocal and nonreciprocal feedback loops with the thalamus, modulating thalamocortical activity through gain control and temporal filtering (Llano and Sherman, 2008; Olsen et al., 2012; Mease et al., 2014; Guo et al., 2017; Clayton et al., 2021; Ibrahim et al., 2021). We observed an overall increase in the intrinsic and synaptic excitability of CTs one day post-NIHL, indicative of an enhanced overall CT excitability (**Figure 3F**). Since the corticothalamic input from CTs to the non-lemniscal thalamic regions is also increased after noise trauma (unpublished results from our lab currently in revision), and the non-lemniscal thalamic regions are thought to be involved in higher-level processing related to internal state and prediction, this increase may reflect a homeostatic plasticity mechanism aimed at enhancing perceptual recovery after NIHL, likely via enhanced contextual modulation (Jones, 1998, 2001; Super et al., 2001; Llinas et al., 2005; Larkum, 2013).

After NIHL, CTs exhibited a persistent reduction in responsiveness to low-intensity sounds (**Figure 6H-K**), which cannot be explained by either increased thalamocortical input or intrinsic excitability. Although the hyperpolarized V_rest_ observed 1 day after NE (**Figure 3B**) might contribute, this is unlikely given that V_rest_ recovers 7 days after NE, but responsiveness to low-intensity sounds does not. This suggests that the observed plasticity in response properties may involve changes in intracortical inputs. The reduction in low-intensity sound responsiveness may indicate long-term changes in the ability to detect quiet sounds, possibly reflecting a trade-off between sound detection and discrimination capabilities after NIHL, a function that has been prescribed to corticothalamic circuitry (Guo et al., 2017). This aligns with the role of CTs in modulating the temporal dynamics and gain of thalamocortical responses, potentially prioritizing detection of more reliable auditory signals.

In contrast to CTs, ETs did not show any changes in their intrinsic excitability after NIHL (**Figure 5**), consistent with previous studies (Nogueira et al., 2023). Although we did not observe any changes in *q* in TC→ET synapses (**Figure 4A-D**), which is the safest metric based on the limitations of our approach, we observed reduced thalamocortical input to ETs one day after noise exposure (**Figure 4E-F_1_**), perhaps due to a decrease in *n* or to the limitations of our approach. However, this reduction is also consistent with the robust shift in the TC input from equivalent between CT and ET to CT dominant one day after NE.

When we explored the sound response properties of ETs, we observed a transient shift in their tuning properties following NIHL. Namely, one day after NE, ETs exhibited a reduction in responses to higher-frequency tones, accompanied by an increase in responsiveness to lower frequencies (**Figure 7**). This reduction in response is consistent with the hypothesized reduced TC input to ETs one day after NE (**Figure 4E-F_1_**). The subsequent recovery of frequency tuning by seven days after NE is also consistent with the recovery of the lemniscal TC input to ETs (**Figure 4E,F_2_**), and suggests that ACtx can dynamically recalibrate itself to restore normal auditory processing capabilities following transient disruptions.

Given our findings on the minimal plasticity changes in ETs after NIHL, the shift in responsiveness of L5 ETs to lower frequencies is likely due to enhanced excitatory cortical inputs. This hypothesized enhanced cortical input coincides with observed increased activity of the ET input to the inferior colliculus after noise trauma (Asokan et al., 2018).

Our study complements and extends the demonstrated cell-type-specific plasticity among ACtx L2/3 interneurons post NE (Kumar et al., 2023). This pattern of plasticity, characterized by differential recovery of inhibitory neuron subtypes in L2/3, highlights the distinct adaptive strategies employed across neurons and ACtx layers, whereby cell-type-specific plasticity in superficial layers may prioritize gain control and network stability, whereas cell-type-specific plasticity in deep layers may engage in adaptive mechanisms that preserve the balance between detection/discrimination and higher-order processing functions (Guo et al., 2017). Although these mechanisms are involved in homeostatic plasticity aimed to preserve hearing after cochlear damage, the same mechanisms can also be involved in the generation of tinnitus and/or hyperacusis, which are associated with noise-induced plasticity across all auditory brain centers, including auditory cortex (Henton and Tzounopoulos, 2021).

In conclusion, our work reveals cell-type-specific plasticity in ACtx ETs and CTs following noise trauma. The observed changes in synaptic strength, intrinsic properties, and sound response tuning suggest a coordinated adaptive response aimed at preserving auditory function despite significant peripheral damage. These findings contribute to a broader understanding of cortical plasticity in sensory processing and may inform therapeutic strategies for mitigating the effects of hearing loss and hearing loss-related disorders, such as tinnitus and hyperacusis.

## Funding

This work was supported by NIH grants R01-DC019618 (TT), R01-DC020923 (TT), R21-DC018327 (RSW), R01-DC020459 (RSW), a Hearing Health Foundation Emerging Research Grant (RSW), and a Klingenstein-Simons Fellowship in Neuroscience (RSW).

## Acknowledgements

We thank current and former members of the Tzounopoulos and Williamson Labs for helpful feedback and discussions, and assistance with animal care.

## Author Contributions

Conceptualization: TT/RSW/SK

Supervision: TT/RSW

Methodology: YZ/SK/NAS/SL/MPA

Formal Analysis: YZ/SK/NAS/RFK

Writing – Original Draft: TT/RSW/SK

Writing – Review and Editing: YZ/SK/NAS/TT/RSW/MPA

## Figure Legends

**Figure S1.**
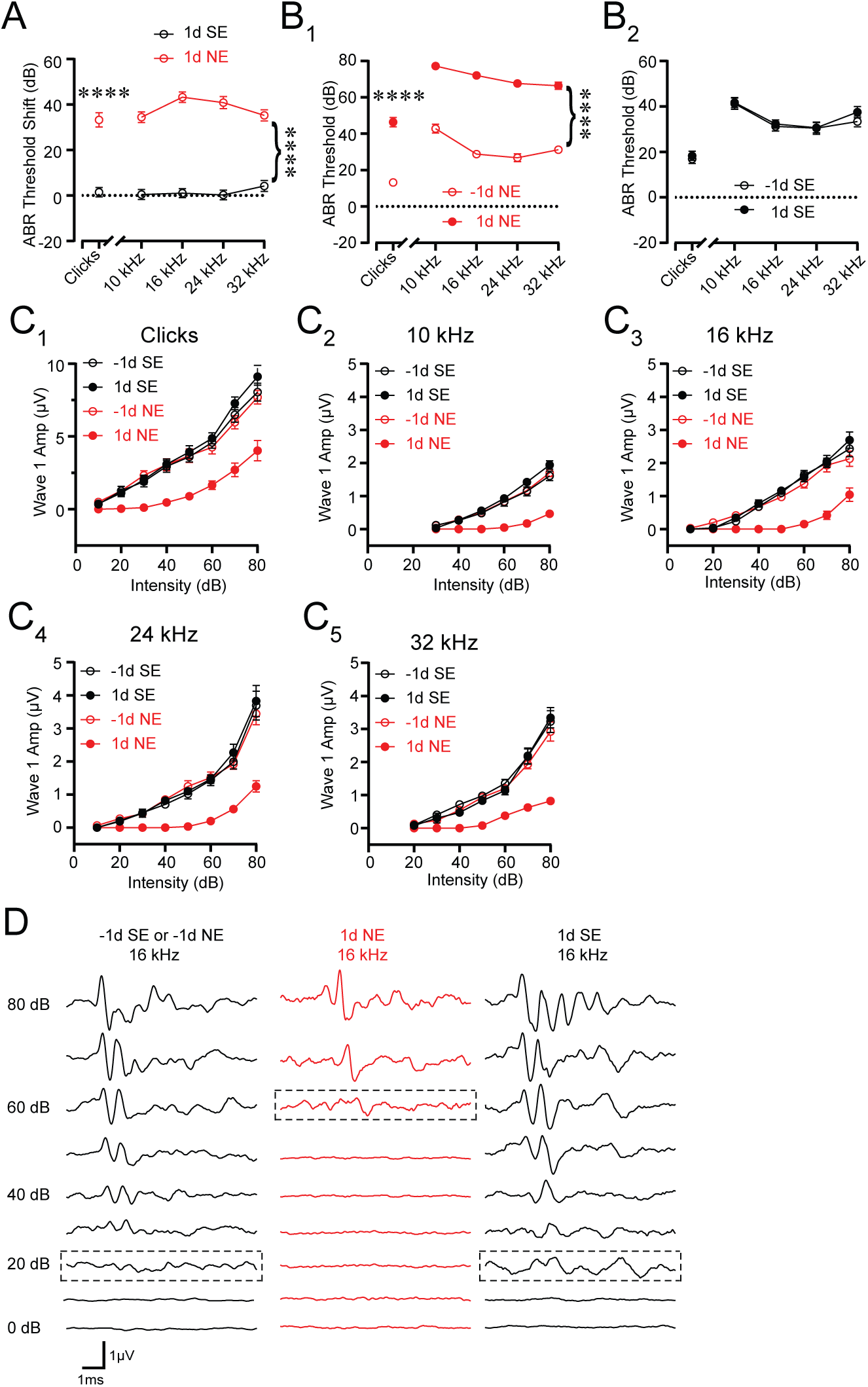
Changes in ABR threshold and Wave I amplitude 1 day after NE. **(A)** ABR threshold shifts 1d after SE and NE. For clicks and different tone frequencies, 1d SE: 24 mice; 1d NE: 25 mice; Asterisks indicate significant differences (****p<0.0001, clicks: Mann-Whitney test; frequency: two-way ANOVA with Bonferroni correction for multiple comparison). **(B)** ABR absolute thresholds for 1d NE (B1) and 1d SE (B2). For clicks and different tone frequencies, -1d NE: 25 mice; 1d NE: 25 mice; -1dSE: 24 mice; 1d SE: 24 mice; Asterisks indicate significant differences (****p<0.0001, clicks: Mann-Whitney test, frequency: two-way ANOVA with Bonferroni correction for multiple comparison). **(C)** Wave I amplitudes for -1d SE, 1d SE, -1d NE and 1d NE mice for clicks (C_1_), 10 kHz (C_2_), 16 kHz (C_3_), 24 kHz (C_4_) and 32 kHz (C_5_). -1d SE: 14 mice; 1d SE: 14 mice; -1d NE: 14 mice; 1d NE: 14 mice. Detailed statistical values are listed in tables. **(D)** Representative traces of ABR recordings (in response to 16 kHz tones) with thresholds highlighted (dotted rectangle) before (left), 1d NE (middle), and 1d SE (right). Detailed statistical values are listed in Table 2.

**Figure S2.**
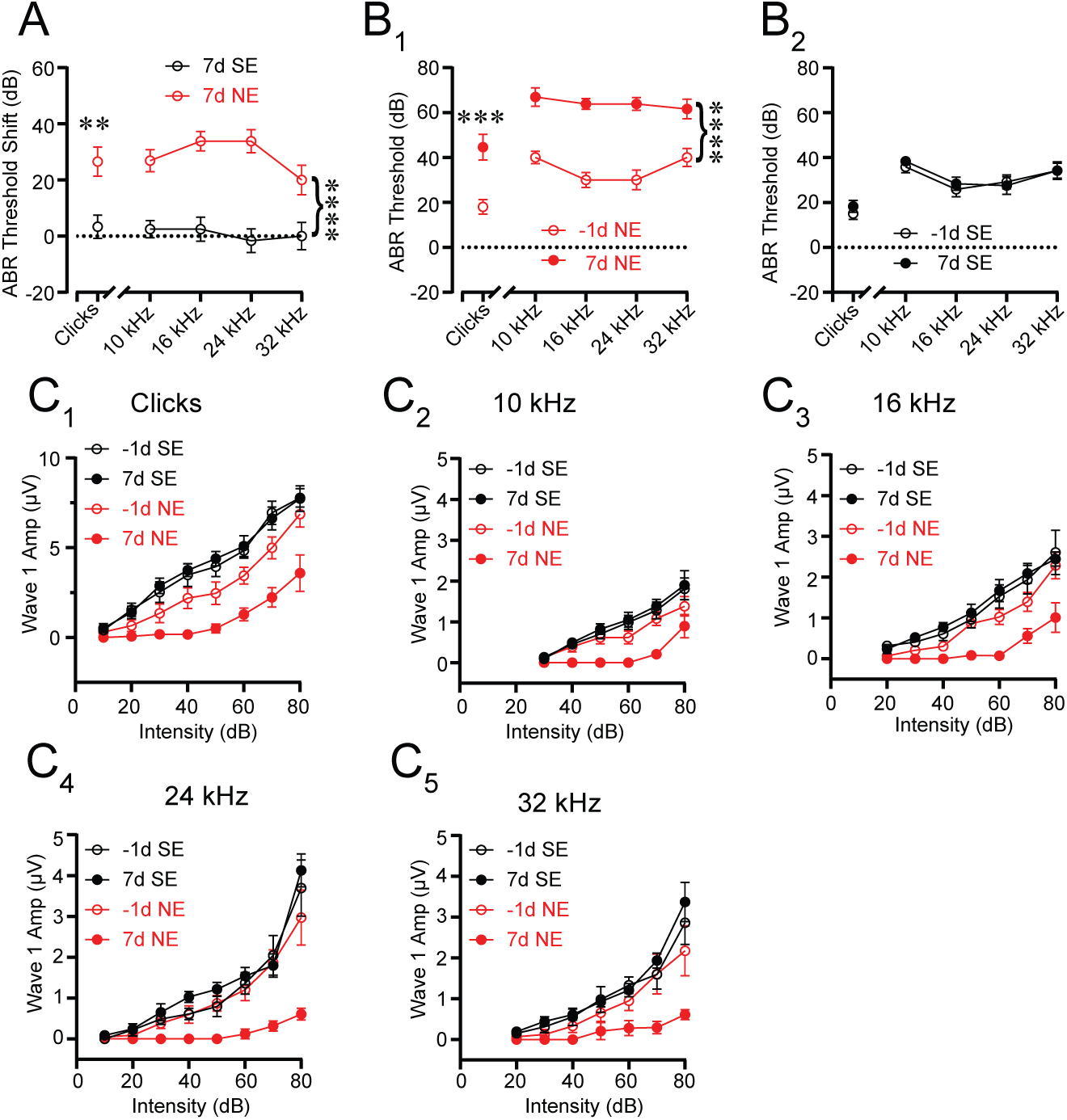
Changes in ABR threshold and Wave I amplitude do not recover 7 days after NE. **(A)** ABR threshold shifts 7d after SE and NE. For clicks and different tone frequencies, 7d SE: 12 mice; 7d NE: 13 mice; Asterisks indicate significant differences (**p<0.01, clicks, Mann-Whitney test; ****p<0.0001, frequency, two-way ANOVA with Bonferroni correction for multiple comparison). **(B)** ABR absolute thresholds in 7d NE (B_1_) and 7d SE (B_2_). For clicks and frequency, -1d NE: 13 mice; 7d NE: 13 mice; -1d SE: 12 mice; 7d SE: 12 mice; Asterisks indicate significant differences (***p<0.001, clicks, Mann-Whitney test; ****p<0.0001, frequency, mixed-effects with Bonferroni correction for multiple comparison). **(C)** Wave I amplitudes between -1d -SE, 7d SE, -1d -NE and 7d NE for clicks (C_1_), 10 kHz (C_2_), 16 kHz (C_3_), 24 kHz (C_4_) and 32 kHz (C_5_). -1d -SE: 6 mice; 1d SE: 6 mice; -1d NE: 6 mice; 7d NE: 6 mice. Detailed statistical values are listed in Table 2.

## Notes

### Competing Interest Statement

The authors have declared no competing interest.

### Summary of Updates

We upload this version, so that we post the original submission. The main difference is that this version lacks Figure 8. Moreover, the previous version has more discussion dedicated to the linking of our in vitro and in vivo findings

